# A cell separation checkpoint that enforces the proper order of late cytokinetic events

**DOI:** 10.1101/308411

**Authors:** Jennifer L. Brace, Matthew D. Doerfler, Eric L. Weiss

## Abstract

Eukaryotic cell division requires sequence dependency relationships in which late processes commence only after early ones are appropriately completed. We have discovered a system that blocks late events of cytokinesis until early ones are successfully accomplished. In budding yeast, cytokinetic actomyosin ring contraction and membrane ingression are coupled with deposition of an extracellular septum that is destroyed immediately after its completion by secreted enzymes. We find this secretion event is linked to septum completion and forestalled when the process is slowed. Delay of septum destruction requires Fir1, an intrinsically disordered protein localized to the cytokinesis site that is degraded upon septum completion but stabilized when septation is aberrant. Fir1 appears to protect cytokinesis in part by inhibiting a separation-specific exocytosis function of the NDR/LATS kinase Cbk1, a key component of a “hippo” signaling pathway that induces mother-daughter separation. We term this system “enforcement of cytokinesis order” (ECO), a checkpoint ensuring proper temporal sequence of mechanistically incompatible processes of cytokinesis.

## Introduction

Eukaryotic cells reproduce through interlaced, mechanistically diverse events that happen with specific relative timing. This sequential order can be crucial: in some cases productive division requires dependency relationships in which late events are not initiated until specific early processes are fully completed, even though these late events do not inherently require the early ones (Hartwell, 1971). Anaphase separation of chromosomes, for example, must not begin until DNA replication is complete and kinetochores are appropriately attached to the mitotic spindle, and cells must not physically divide before the duplicated genome has been partitioned. The ancient cyclin:CDK regulatory system creates order by inducing, preventing, or permitting semi-autonomous processes of cell division (Oikonomou and Cross, 2010; Sullivan and Morgan, 2007). For example, key early steps in formation of DNA replication origins are only possible at low CDK activity, while a single initiation of DNA replication from these origins can only occur at high CDK activity (reviewed in Siddiqui et al., 2013). In a different case, an inherently oscillatory process that releases and re-sequesters a key mitotic phosphatase initiates only at high mitotic CDK activity, effectively locking the phosphatase release oscillator to a specific phase of the CDK activity cycle (Lu and Cross, 2010; Manzoni et al., 2010).

In addition to linking stages of division to the cyclin-CDK system, eukaryotic cells have evolved regulatory mechanisms known as checkpoints that actively block downstream events until upstream ones are successfully finished (Hartwell and Weinert, 1989; Khodjakov and Rieder, 2009; Li and Murray, 1991). These systems effectively monitor the status of key processes, generating negative signals that impede progression to later stages until specific biochemically-sensed criteria are satisfied. Checkpoint signaling can function by impinging on the cyclin:CDK system itself: for example, unattached kinetochores produce an inhibitor that blocks destruction of mitotic cyclin and thus prevents the metaphase-anaphase transition (reviewed in Musacchio, 2015). Importantly, checkpoint-monitored processes lose sequential dependencies when checkpoint mechanisms are nonfunctional, a disruption that is especially problematic when normally early events are themselves disrupted. The spindle assembly and DNA damage checkpoints are well studied, and it is becoming clear that additional checkpoint-like mechanisms protect the integrity of cell division. For example, failure to successfully complete cytokinesis in higher eukaryotes induces a checkpoint-like response that prevents tetraploidization (Steigemann et al., 2009) and cells with lagging chromosomes actively block cytokinesis that would cause chromosome damage (Mendoza et al., 2009; Nähse et al., 2017; Norden et al., 2006).

Eukaryotic cells undergo dramatic reorganization at the end of mitosis, producing two cells from one through the processes of cytokinesis. This division requires execution of mechanistically diverse events in an unvarying and often rapid sequence, in which late processes are not initiated until early ones are effectively completed. Specification of the division site and assembly of cytokinetic structures precedes the mechanical and regulatory events of actomyosin ring (AMR) constriction and membrane ingression; this is followed by disassembly of cytokinetic machinery, cessation of cytokinetic membrane trafficking, and abscission of the divided cells (Glotzer, 2016; Gould, 2015; Green et al., 2012; Mierzwa and Gerlich, 2014). In some cases the relative timing of cytokinesis events may reflect inherent structural dependencies, but overall the mechanisms that enforce temporal sequence of cytokinesis phases are not well understood.

Cytokinesis in the budding yeast *Saccharomyces cerevisiae* proceeds through a rapid sequence of processes that are broadly conserved (reviewed in Balasubramanian et al., 2004; Bhavsar-Jog and Bi, 2017; Juanes and Piatti, 2016; Weiss, 2012), including AMR construction and constriction, highly localized membrane addition, and membrane abscission. Like many eukaryotes, budding yeast cells build a specialized extracellular barrier called the septum at the site of cytokinesis: in general, free-living cells like budding yeast are under extreme turgor pressure, and the septum is thus critical for osmotic integrity during the division process (Cortés et al., 2012; Levin, 2005; Proctor et al., 2012). Primary septum (PS) construction in budding yeast structure is intimately associated with actomyosin ring (AMR) constriction, with the transmembrane enzyme Chs2 acting specifically at the site of membrane ingression to locally synthesize and extrude chitin polymer (Figure 1A i) (Devrekanli et al., 2012; Nishihama et al., 2009; Schmidt et al., 2002; Shaw et al., 1991).

**Figure 1:**
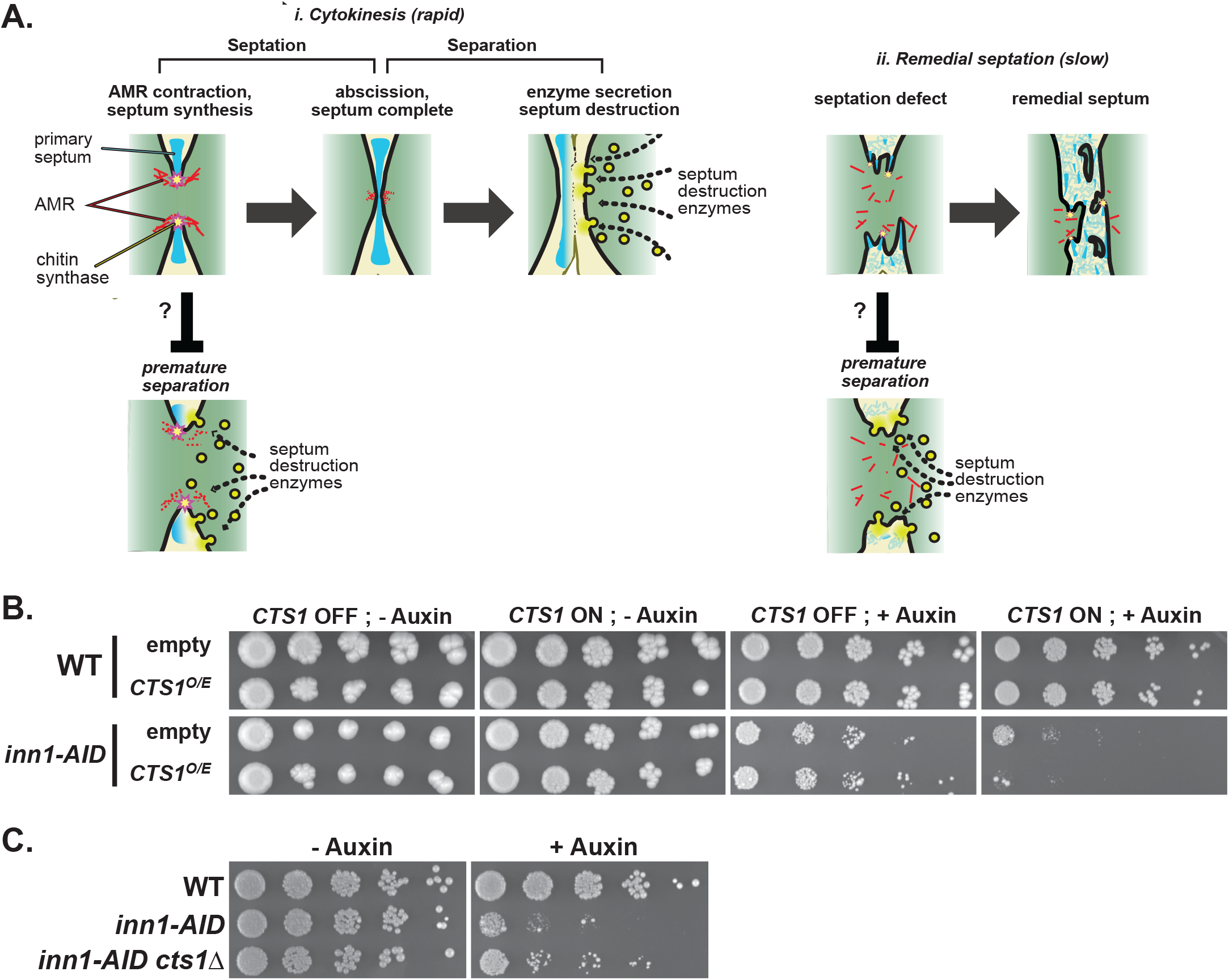
Septation mutants are sensitive to inappropriate activation of cell separation. (A) Cytokinesis in budding yeast. (i) Normal, rapid cytokinesis can be separated into two broad phases: “septation” and “cell separation”. Only after completion of septation are septum destroying enzymes secreted. To ensure this temporal order, we predict cells actively inhibit separation until septation is complete. (ii) Cells with septation defects generate a remedial septum. Cells forming a slow remedial septum are particularly sensitive to premature septum destruction, and we propose cells activate a checkpoint-like mechanism to enforce the strict temporal order of septation and separation. (B) Inappropriate *CTS1* expression is detrimental when septation fails. Wild type (WT) and *inn1-AID* cells transformed with an empty vector or a galactose inducible vector expressing *CTS1* were spotted in five-fold serial dilutions to plates containing glucose (*CTS1* OFF) or galactose (*CTS1* ON). Additionally, the plates contained 0.5 mM auxin (+ auxin) or DMSO (-auxin). Plates were incubated at 30° C for 3 days. All strains express the E3 ligase *TIR1*. (C) *CTS1* deletion restores viability to cells with disrupted septum synthesis. The indicated strains were grown on YPD plates containing 0.5 mM auxin (+ auxin) or DMSO (−auxin) and incubated at 30° C for 3 days. All strains express the E3 ligase *TIR1*.

Both AMR constriction and septum synthesis are triggered in budding yeast by the mitotic exit network (MEN) (reveiwed in Meitinger et al., 2012), a conserved ‘hippo’ signaling pathway activated as dividing cells pass the metaphase-to-anaphase transition, as well as by localized activation of the GTPase Rho1 (Onishi et al., 2013; Yoshida et al., 2006, 2009). Exemplifying complex multi-system coordination needed for successful cytokinesis, MEN activation releases the membrane-spanning chitin synthase Chs2 from sequestration in the ER, allowing its trafficking to the cytokinesis site, and induces AMR localization of the proteins that activate Chs2 at the ingressing division furrow (Chin et al., 2012; Kuilman et al., 2015; Meitinger et al., 2010; Nishihama et al., 2009; Oh et al., 2012; Palani et al., 2012; Verplank and Li, 2005; Zhang et al., 2006). As the AMR-guided chitin septum forms, it is followed by localized production of glucan-rich secondary septa (SS) on both mother and daughter cell sides (Cabib, 2004; Lesage and Bussey, 2006). Formation of the SS involves activation of the glucan synthase Fks1, the exocyst component Sec3, and the chitin synthase Chs3 by Rho1 (Yoshida et al., 2009). While mechanisms that coordinate timing of PS and SS production are incompletely understood, recent analyses demonstrate that factors promoting early events can directly inhibit later events, including regulation of the GTPases Rho1 and Cdc42 and the chitin synthase Chs3, to help ensure appropriate order of these distinct processes (Atkins et al., 2013; Meitinger et al., 2013; Oh et al., 2017; Onishi et al., 2013).

Remarkably, the septum is destroyed only minutes after it is completed. Once septation is complete, daughter cells secrete degradative enzymes to destroy the primary septum and release daughter cells from their mothers (Figure 1A i). A ‘hippo’ signaling pathway called the RAM network (<u>R</u>egulation of <u>A</u>ce2 and <u>M</u>orphogenesis) controls this process (reviewed in Weiss, 2012). Mutants in this pathway exhibit defects in cell separation and accumulate as large clumps of cells (Kurischko et al., 2005; Nelson et al., 2003; Weiss et al., 2002). The downstream most kinase, Cbk1, is critical to direct the cell separation process (Bidlingmaier et al., 2001; Colman-Lerner et al., 2001; Jansen et al., 2006; Weiss et al., 2002). In late mitosis, Cbk1 is activated upon MEN-dependent release of the phosphatase Cdc14 and activates the transcription factor Ace2 (Brace et al., 2011). Cbk1 interacts with Ace2 through a recently discovered docking motif (Gógl et al., 2015; Nguyen Ba et al., 2012) and phosphorylates the transcription factor Ace2 at its nuclear export sequence trapping the protein in the daughter cell nucleus (Mazanka et al., 2008). There, Ace2 drives transcription of genes required to efficiently degrade the septum (Colman-Lerner et al., 2001). One such gene encodes the endochitinase Cts1, a secreted protein that degrades chitin in the primary septum leading to cell separation (Elango et al., 1982; Kuranda and Robbins, 1991); however, any regulation on Cts1 secretion is yet unknown.

It is unclear how - or if - synthesis of the septum is coupled to its destruction. Interestingly, Ace2 localizes to the daughter cell nucleus prior to the completion of septation, and transcription of Cts1 and other cell separation genes begins prior to cytokinesis (Mazanka et al., 2008). Thus, cells could be capable of producing Cts1 and other degradative enzymes, even though premature destruction of the forming structure would likely be detrimental to the dividing cell (Cabib et al., 1992). Thus, we hypothesized that a checkpoint mechanism forestalls separation processes until completion of septation (Figure 1A). To better understand mechanisms that enforce cytokinesis order, we investigated the coupling of septation and mother-daughter separation in yeast cytokinesis. Our findings indicate that a checkpoint mechanism delays secretion of Cts1, and thus destruction of the septum, when early cytokinetic processes important for septum formation are defective. This pathway involves the protein Fir1, a probable RAM network target that is normally degraded after AMR constriction but persists at the cytokinesis site when septation is defective. In the absence of Fir1, cells with septation defects prematurely secrete Cts1 and fail to complete cytokinesis normally. We term this checkpoint-like mechanism the “Enforcement of Cytokinesis Order” pathway (ECO).

## Results

For clarity, we refer to the combined processes of AMR contraction, membrane ingression, septum construction, abscission, and septum degradation as “cytokinesis” (Figure 1A). We divide cytokinesis into two broad phases: “septation” beginning with AMR contraction / septum construction and ending with septum completion, and “separation” beginning with secretion of Cts1 and other enzymes and ending with full release of the extracellular mother-daughter junction (Figure 1A i). In budding yeast, these events happen in rapid succession, leading to production of two completely separated cells in about 10 minutes under normal growth conditions.

### Cells with septation defects are sensitive to artificial changes in Cts1 levels

We sought to determine if cells enforce dependency of late stages of cytokinesis on successful completion of early ones. The protein Inn1 links the actomyosin ring to machinery that extrudes chitin polymer into the extracellular space (Devrekanli et al., 2012; Meitinger et al., 2010; Sanchez-Diaz et al., 2008), and is crucial for successful early cytokinesis. Cells lacking Inn1 cannot perform actomyosin-directed septum formation, and instead deposit a disorganized remedial septum composed of chitin and other extracellular glucans at the bud neck that eventually closes off the cytoplasmic connection between mother and daughter cells (Nishihama et al., 2009, Figure 1A ii). We used a system in which the exogenous ubiquitin ligase Tir1 polyubiquitinates proteins carrying an auxin-inducible degron (AID) tag upon auxin addition (Nishimura et al., 2009) to conditionally deplete Inn1 and disrupt normal septation. We found that, as reported, cells carrying the *inn1-AID* allele produce a functional Inn1-AID fusion protein that is rapidly degraded in the presence of auxin, with attendant failure of mother-daughter separation (Figure S1A, 1B and (Devrekanli et al., 2012; Nishihama et al., 2009; Sanchez-Diaz et al., 2008)).

Complete failure of cytokinesis is detrimental to budding yeast, and cells with septum formation problems exhibit poor viability (Korinek et al., 2000; Luca et al., 2001; Nishihama et al., 2009; Tolliday et al., 2003; Vallen et al., 2000). The processes of mother-daughter separation are directly antagonistic to septation. If these processes do not inherently require septum completion to proceed, then enhancing their activities might worsen phenotypes associated with septation defects. Conversely, elimination of cell separation might ameliorate them. We therefore overexpressed the cell separation chitinase Cts1 in cells with defective early cytokinetic processes, placing *CTS1* under the control of a galactose-inducible promoter and again using auxin treatment of *inn1-AID* cells induce defective septation. We found that *inn1-AID* cells grew poorly but measurably in the presence of auxin under induction conditions, while no growth occurred in combination with the *CTS1* overexpression vector (Figure 1B; compare empty vector to *CTS1*^O/E^, *CTS1* ON, +Auxin). Notably, this enhanced lethality was only seen in combination with failed septation: wild type cells or *inn1-AID* cells in the absence of auxin were not appreciably sensitive to *CTS1* overexpression. In contrast, we found that deleting *CTS1* partially rescued the poor growth of *inn1-AID* strains on auxin-containing media (Figure 1C).

### Cts1 secretion is blocked in cells with cytokinetic defects

Given that overexpression or absence of Cts1 alters the phenotypic effect of early cytokinesis defects, we sought to determine if the action of this key enzyme in mother-daughter separation is delayed when septation is defective. The RAM network ‘hippo’ signaling system activates transcription of the *CTS1* gene at the M/G1 transition and relieves translational repression of the *CTS1* mRNA (Dohrmann et al., 1992; Jansen et al., 2009). Additionally, Cts1 is a secreted protein that transits the endomembrane system before secretion to the extracellular space (Correa et al., 1982; Elango et al., 1982). Thus, we considered multiple levels at which the function of Cts1 could be regulated in response to septation failure, including *CTS1* gene transcription, *CTS1* mRNA translation, and the trafficking itinerary of Cts1 prior to secretion. The *CTS1* mRNA 3’ and 5’ UTRs carry important though incompletely defined regulatory elements (Aulds et al., 2012; Jansen et al., 2009; Wanless et al., 2014). Since N- or C-terminal tagging could disrupt this regulation, we generated a strain expressing Cts1 with an HA-tag internal to the protein. Since *CTS1* expression peaks only briefly following mitotic exit (Brace et al., 2011; Mazanka et al., 2008), we examined *CTS1* mRNA, protein expression, and secretion in a synchronized population of cells undergoing a single cytokinetic event.

Auxin treatment of Tir1-expressing wild type cells did not alter the kinetics of cell cycle progression, *CTS1* transcription, translation, glycosylation and/or secretion (Figure S1C - E). Cells were synchronized in metaphase, treated with auxin or vehicle and then released from the arrest in the presence or absence of auxin. We collected cell samples through a single round of cell division were processed for RNA, protein and budding index analysis. We saw no difference in budding index (Figure S1E), *CTS1* transcription (Figure S1D), translation or secretion (Figure S1C) in the presence or absence of auxin. Next, we examined Cts1’s itinerary during a single cell division event in which cytokinesis was disrupted. Use of the *inn1-AID* genotype allowed us to directly compare the same population of cells with normal (minus auxin) or failed septation (plus auxin). We observed identical induction of *CTS1* transcription following mitotic exit in cells treated with or without auxin, 0 – 60 minutes following release (Figure 2C), demonstrating that regulation occurs downstream of the transcription factor, Ace2.

**Figure 2:**
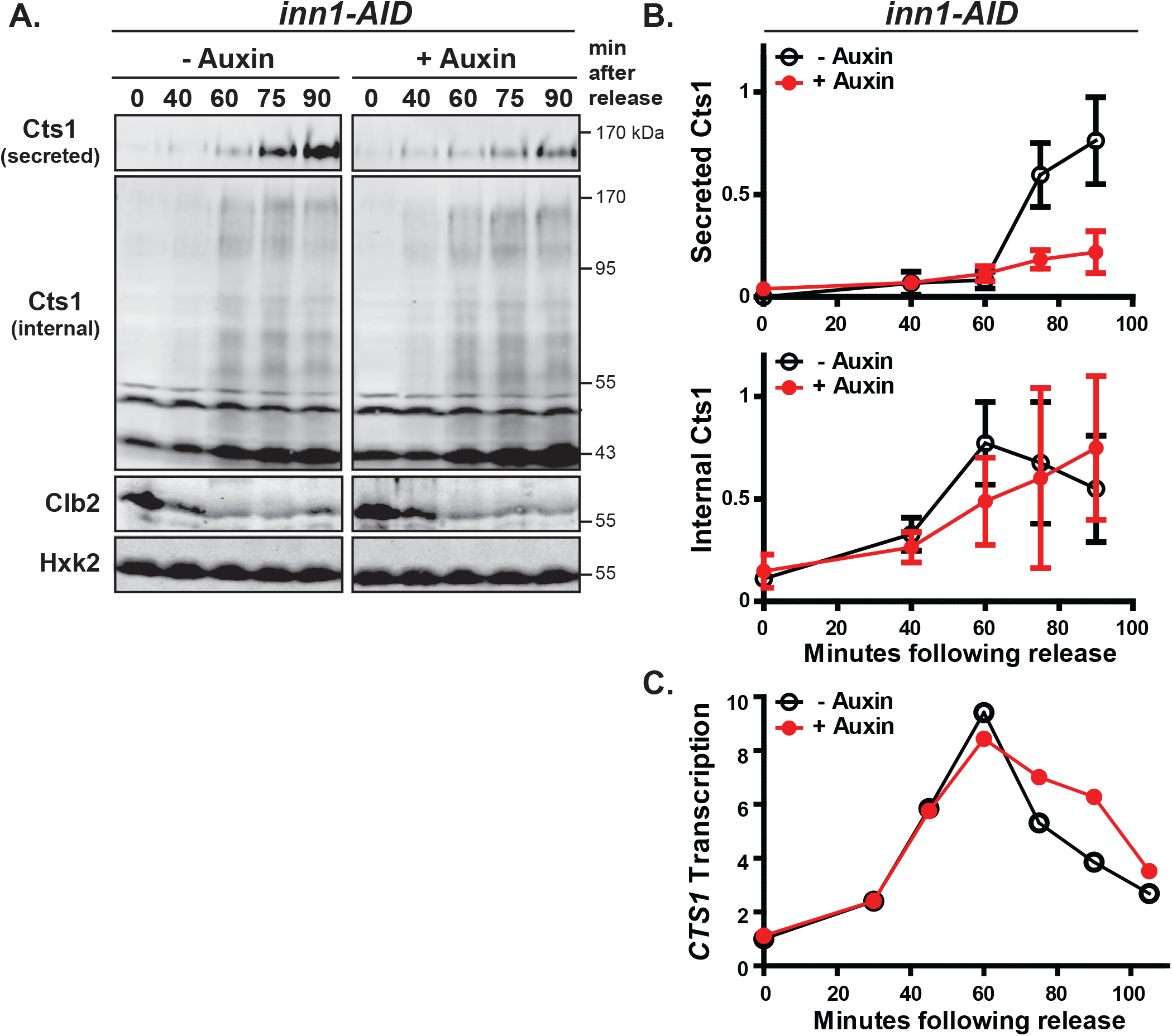
Septation failure is associated with a block in secretion of the septum degrading enzyme, Cts1. (A) Cts1 secretion is blocked when septation fails. *inn1-AID* cells expressing HA-tagged Cts1 were synchronized in mitosis and treated with DMSO (-auxin) or 0.5 mM auxin (+ auxin). Protein was collected at the indicated times following mitotic release at 25° C. A Western blot of secreted Cts1 (top panel) and internal pool of Cts1 (middle panel) is shown. The asterisk indicates a non-specific band (see Figure S1C). Hexokinase (Hxk2) and the cyclin Clb2 are shown as a loading and cell cycle release control, respectively (lower panels). A representative blot is shown. (See also Figure S1). (B) Western blot quantification. Normalized signal of secreted Cts1 (upper panel) and internal Cts1 (lower panel) is shown. Error bars represent the standard deviation from three independent time courses. (C) Cts1 transcript induction is not altered upon failed septation. *CTS1* mRNA from cells in (A) were measured by qPCR at the time indicated after release. The fold change in *CTS1* mRNA (normalized to *ACT1*) relative to minus auxin at time 0 is shown. A representative time course is shown.

We then examined Cts1 translation and secretion to the media. Cts1 transits the endomembrane system where it is heavily O-mannosylated (Gentzsch and Tanner, 1997; Kuranda and Robbins, 1991). Cell lysate (internal fraction) includes non-secreted Cts1 found in the ER, Golgi and secretory vesicles and runs as a smear of different glycosylation forms. Cts1 isolated from the media (secreted fraction), runs as the most heavily glycosylated, mature form (Figure 2A). We collected cells following metaphase arrest and release and analyzed internal and secreted fractions of Cts1 qualitatively (Figure 2A) and quantitatively (Figure 2B). During a normal cell division (minus auxin), Cts1 (internal) accumulated 40 minutes following release, peaking at 60 minutes. After peak internal protein production, secreted Cts1 accumulated in the media (75 minutes) (Figure 2A and B).

Under conditions of failed septation (plus auxin), the level of internal Cts1 mirrored that of cells undergoing normal cytokinesis (Figure 2A and B). Additionally, we saw no obvious change in the glycosylation pattern of internal Cts1 (Figure 2A). These data suggest Cts1 co-translational insertion of into the ER and glycosylation are not grossly affected upon failed septation. More strikingly, however, we saw a significant delay in the timing and amount of Cts1 secreted to the media in cells lacking Inn1. When cytokinesis proceeded normally we detected robust Cts1 secretion into the media at 75 minutes, while finding very little secreted Cts1 at the same time in cells undergoing defective septation (Figure 2A and B). Even at a later time point, very little Cts1 was secreted. To confirm this secretion block was due to depleted Inn1 and not auxin treatment we examined Cts1 secretion in auxin treated *inn1-AID* cells lacking the E3 ligase Tir1 (required for AID-tagged protein degradation) and saw the Cts1 secretion pattern was similar to wild type cells and minus auxin samples (Figure S1F).

To corroborate these results we ran a similar experiment in cells expressing an untagged version of Cts1. Using an antibody against the endogenous protein, we found that Cts1 secreted to the media was reduced in cells depleted of Inn1 (Figure S1G). This antibody, unfortunately, was unable to detect the internal fraction of Cts1 (data not shown). Additionally, we measured enzymatic activity of chitinase secreted to the media and found that cells lacking Inn1 exhibited reduced chitinase activity in the cell associated/media fraction compared to control cells (Figure S1H). In sum, our findings indicate that transcription, translation and glycosylation of Cts1 are not grossly affected upon failed septation, but that a significant block in Cts1 secretion occurs.

### Identification of a regulatory component monitoring cytokinesis

Our results suggest that cells detect septation defects and block the subsequent step, secretion of chitinase, to ensure the proper order of cell division events. Identification of factors that guarantee the dependency of the system would provide additional evidence of a regulatory system. Cells can rely on an inherent dependency: physically septum synthesis and destruction cannot occur at the same time, or it could rely on enforcement of a regulatory block to prevent separation until septation is complete. Chitinase overexpression killed cells with disrupted septation suggesting these processes can occur at the same time. Consequently, we propose the mechanism involves enforcement of a regulatory block and expected elimination of this regulation might cause premature septum destruction and lethality. Thus, we examined high-throughput SGA (synthetic genetic array) data (Costanzo et al., 2010) to identify gene products demonstrating a negative genetic interaction with known cytokinetic factors. One such gene, *FIR1*, has no known role in cytokinesis yet exhibits negative genetic interactions with several cytokinetic factors, including *GPS1, DBF2, CYK3, FKS1*, and *RGL1*.

Fir1 is largely an unstructured protein with no predicted domains. Interestingly, Fir1 is enriched in short linear motifs (SLiMs): many predicted to dock with cell cycle proteins (Cks1, Cdc5, Cln1/2) or are predicted to be cell cycle dependent modification sites including cyclin dependent kinase phosphorylation (Holt et al., 2009) and Cdc14 dependent dephosphorylation (Kuilman et al., 2015). The protein also contains several destruction box motifs and a SUMO interaction motif (Figure S2A ii). This, together with evidence that *FIR1* transcripts are cell cycle regulated peaking in G2/M (Spellman et al., 1998) and genome-wide studies demonstrating that Fir1 localizes to the site of septation (Huh et al., 2003), provoked us to further investigate Fir1’s role in monitoring the cell division process.

First we verified that elimination of *FIR1* would exacerbate the lethality of cells that fail septation. As expected for a checkpoint regulatory protein (Li and Murray, 1991; Weinert and Hartwell, 1988), *FIR1* deletion was viable and not required for viability under normal growth conditions (Figure 3A). Cells divided normally and exhibited wild type morphology. However, upon failed septation, *FIR1* became essential: the *inn1-AID fir1*Δ genotype was lethal with auxin treatment (Figure 3A) and loss of *FIR1* was lethal with other factors that are implicated in cytokinesis and septation including *CYK3, FKS1, and RGL1* (Figure S2B).

**Figure 3:**
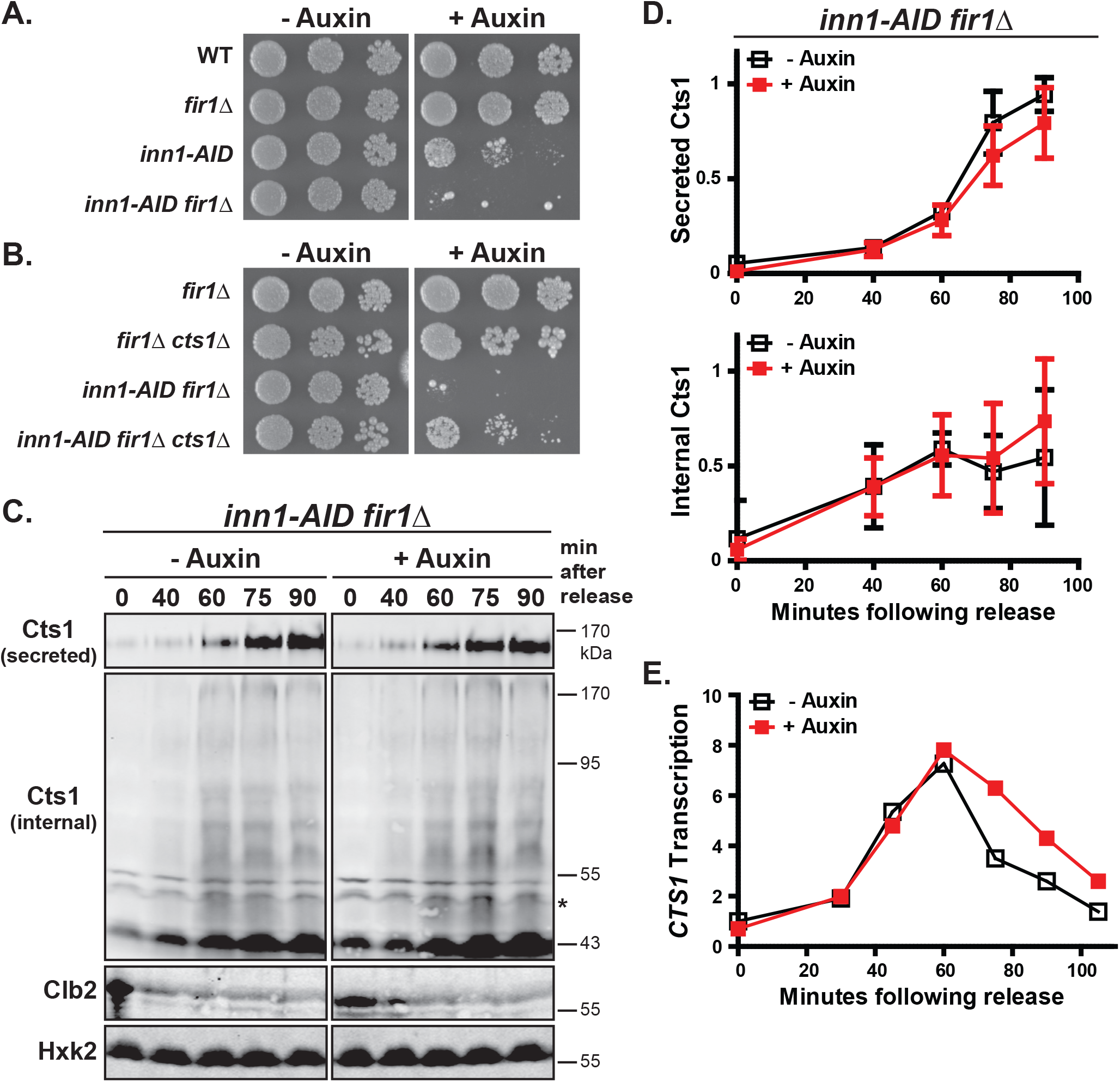
Septation mutant viability requires Fir1 to prevent inappropriate Cts1 secretion. (A) Loss of *FIR1* is detrimental to cells with septation defects. Five-fold serial dilutions of the indicated strains were spotted to YPD media as in Figure 1C. (See also Figure S2B). (B) *CTS1* deletion rescues the lethality of *inn1-AID fir1*Δ. Five-fold serial dilutions of the indicated strains were spotted to YPD as in Figure 1C. (C) Fir1 blocks Cts1 secretion when septation fails. *inn1-AID fir1*Δ cells expressing HA-tagged Cts1 were synchronized in mitosis and treated with DMSO (−auxin) or 0.5 mM auxin (+ auxin). Cells were collected and processed as in Figure 2A. (D) Western blot quantification (as in Figure 2B). (E) *CTS1* transcript induction is not altered by lack of *FIR1. CTS1* mRNA from cells in (C) were measured by qPCR as in Figure 2C. (Figures 3C-E were done in parallel with Figure 2A-C).

Since cells failing to complete septation do not secrete Cts1 and are sensitive to inappropriate expression of *CTS1*, we predicted elimination of a regulatory protein might cause premature septum degradation leading to death. Therefore, if the lethal interaction of *FIR1* and *inn1-AID* is due to premature cell separation, it should be suppressed by *CTS1* deletion. Indeed, *CTS1* deletion restored viability to *inn1-AID fir1*Δ cells treated with auxin (Figure 3B). These data support a role for *FIR1* as a regulatory component of a system to monitor septum completion and suggests its absence leads to bypass of this system.

### Fir1 prevents Cts1 secretion in septation mutants to ensure cell integrity

A block in Cts1 secretion following failure to complete septation (Figure 2), and genetic evidence demonstrating the lethality of *inn1-AID fir1*Δ cells could be reversed by elimination of *CTS1* (Figure 3B), lead us to propose that Fir1 functions to block inappropriate Cts1 secretion. To test this directly, we examined Cts1’s itinerary in *inn1-AID fir1*Δ cells. Concurrently with experiments with *inn1-AID* (Figure 2), *CTS1* transcription, protein production and secretion were analyzed (Figure 3C) and quantified (Figure 3D) in *inn1-AID fir1*Δ cells. Synchronized cells were released into mitosis in the presence or absence of auxin and samples were collected for RNA and protein analysis. We found *FIR1* deletion did not overall alter levels of *CTS1* transcription (compare Figure 2C and 3E), and transcription initiation (0-60 minutes) was similar in cells completing cytokinesis normally (minus auxin) and cells failing to complete septation (plus auxin) (Figure 3E).

Similar to *inn1-AID* cells, *CTS1* translation and glycosylation (internal) were not appreciably altered in the absence of *FIR1* (Figure 3C and D). We saw internal production of Cts1 began around 40 minutes in both plus and minus auxin treated cells and increased over the time course of the experiment. We also did not observe any evidence of changes in overall glycosylation pattern, a similar smearing and banding pattern was seen in the presence or absence of *FIR1* (compare Figure 2A and 3C).

However, we saw a significant difference in Cts1 secretion in cells lacking *FIR1*. Unlike *inn1-AID* cells (Figure 2A and B) which delay Cts1 secretion, auxin treated *inn1-AID fir1*Δ cells secreted Cts1 to the media at levels and timing comparable with mock treated cells (Figure 3C and D). Interestingly, in the absence of auxin, secreted Cts1 levels began to increase at 60 minutes (compared to 75 minutes) in cells lacking *FIR1* compared to wild type cells (compare Figure 2B and 3D, Secreted Cts1) suggesting *FIR1* may also play a slight role in preventing premature Cts1 secretion under normal conditions.

Cells that inappropriately secrete Cts1 should lack chitin in the septum and might fail to complete abscission (Cabib and Schmidt, 2003). We treated synchronized wild type, *fir1*Δ, *inn1-AID*, and *inn1-AID fir1*Δ cells with auxin (or DMSO) and pooled time points to enrich for cells that have just failed or completed cytokinesis. Fixed cells stained with the chitin-binding fluorescent dye calcofluor-white demonstrate the extent of chitin in the bud neck (Pringle, 1991). When septation and cell separation occurred normally (in the absence of auxin or in the presence of auxin where Inn1 is functional), chitin staining was similar in all the genotypes (Figure 4A, -Auxin). However, upon auxin addition, *inn1-AID* cells remained attached at the bud neck with strong chitin staining (Figure 4A, +Auxin, white triangle). This is consistent with evidence that cells forming a remedial septum contain chitin in the septal region generated by the chitin synthase, Chs3 (Cabib and Schmidt, 2003; Shaw et al., 1991). In contrast, *inn1-AID fir1*Δ cells exhibited extensive regions of chitin-free staining between mother and daughter cell (Figure 4A, +Auxin, white arrows). We also noticed several cells forming a new bud through the chitin-free region between cells (asterisk) to form a “zygote-like” morphology. To quantify this morphology, we released *inn1-AID* and *inn1-AID fir1*Δ cells from arrest in the presence or absence of auxin and grew cells to allow the formation of buds. Morphology fell into either a chained or zygote-like morphology, and we found a significant increase in the number of zygote-like cells in the *inn1-AID fir1*Δ strain compared to the *inn1-AID* strain (Figure 4B). We never saw this phenotype in cells not treated with auxin. Consistent with the hypothesis that cells lacking *FIR1* prematurely degrade the septum as it forms, this “zygote-like” morphology was observed in cells that lack bud neck chitin through genetic or chemical means of inhibiting chitin synthases (Cabib and Schmidt, 2003; Shaw et al., 1991).

**Figure 4:**
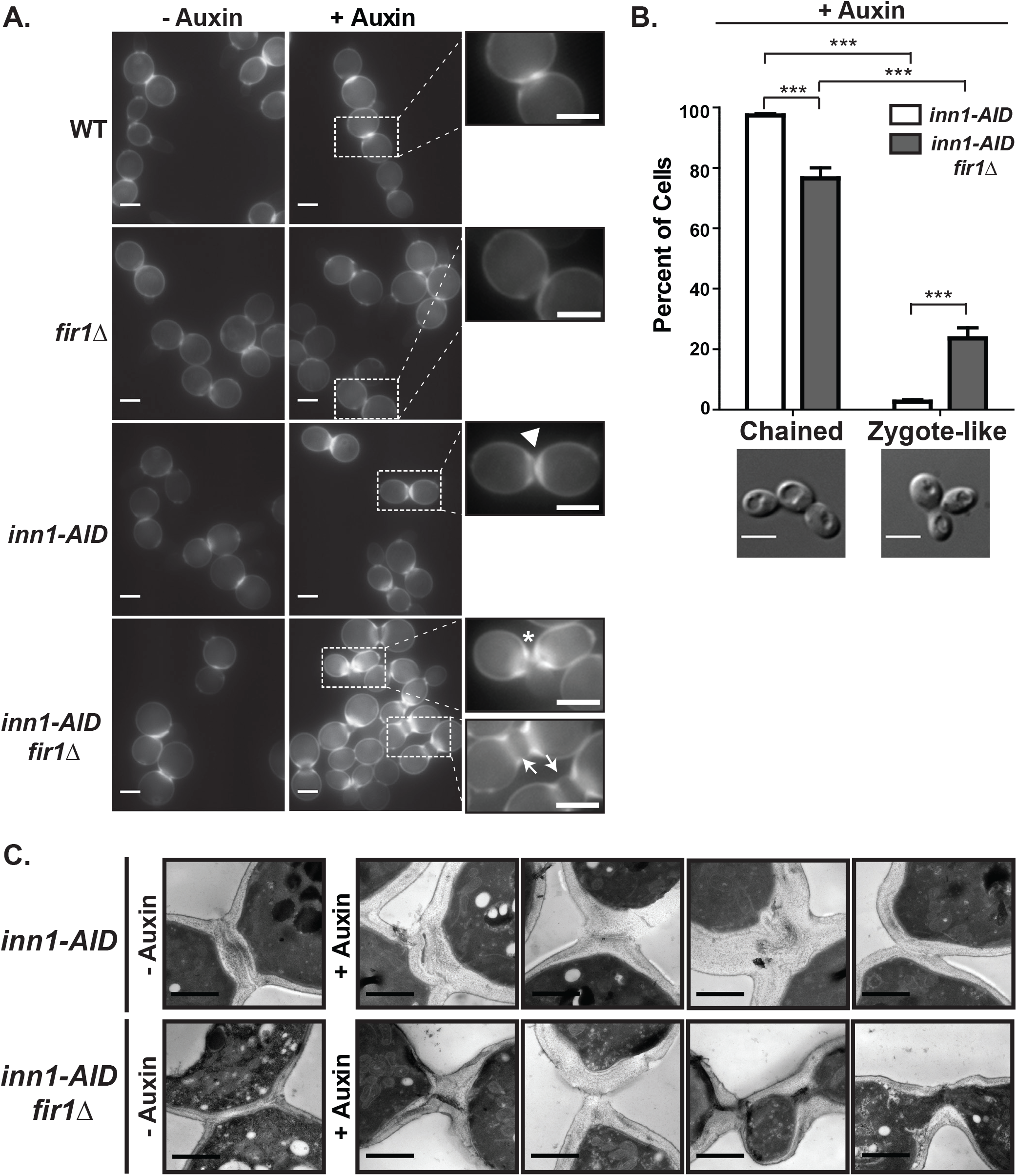
Inappropriate Cts1 secretion causes premature septum destruction leading to failed abscission and thinned septa. (A) Inappropriate Cts1 secretion destroys chitin in the aberrant septum. The indicated genotypes, treated with or without auxin, were synchronized and released at 30° C. Cells were collected at times to enrich for cells completing cytokinesis (see methods). Calcofluor staining of fixed cells demonstrates chitin content. Enlarged region highlights bud neck chitin. Triangle highlights region of increased chitin; arrow decreased chitin; asterisk aberrant bud growth. Representative images are shown. Scale bar is 5 μm. (B) Inappropriate secretion of Cts1 leads to failed abscission and budding defects. Synchronized and auxin treated *inn1-AID* and *inn1-AID fir1*Δ cells were released from arrest at 30° C for 3 hours. Budding morphology was binned into chain-like or zygote-like and the percent of cells exhibiting the indicated morphology (representative image scale bar: 5 μm) is shown. Error bars represent standard deviation of three independent experiments (n>100 each trial); *** represents a p-value < 0.001 (t-test). (C) Cells inappropriately secreting Cts1 fail to complete septation or have thinned septal regions. Synchronized *inn1-AID* and *inn1-AID fir1*Δ cells were treated as in (A) and were processed for electron microscopy. Representative images of each genotype are shown (see also Figure S3). Scale bar represents 1 μm.

Finally, we used electron microscopy to examine the organization of the septum. Synchronized *inn1-AID* and *inn1-AID fir1*Δ cells treated with DMSO or auxin were released from arrest, collected and pooled to enrich for cells completing or failing cytokinesis, respectively. As previously demonstrated (Nishihama et al., 2009), cells lacking *INN1* construct a disorganized, thick remedial septum (Figure 4C and S3, +Auxin). In the additional absence of *FIR1*, we observed thinned remedial septa and found many cells failed to complete abscission upon auxin treatment (Figure 4C and S3, +Auxin). Taken together, these data provide additional evidence that upon septation failure, Fir1 functions to prevent premature septum degradation.

### Cytokinetic defective cells prevent Fir1 degradation

Fir1 localizes to the bud neck (Huh et al., 2003), but its timing and dynamics during cytokinesis are not known. To examine this, we localized Fir1-GFP relative to the actomyosin ring (AMR) component Myo1-mcherry. Myo1 marks the site of septation and contracts and disassembles upon completion of septation (Bi et al., 1998). Fir1 initially co-localized with Myo1 in a ring and filled in a disc-like structure behind the contracting AMR (Figure 5A and Movie S1). Interestingly, it remained momentarily at the bud neck, before its own disappearance. In a population of synchronized cells, we found a similar result: Fir1 and Myo1 were bud neck localized in most cells upon early mitotic release (0-20 minutes), followed by a brief period in which Fir1 was present in more cells than Myo1 (30-45 minutes), and then neither protein was observed at the bud neck in most cells (45-55 minutes) (Figure 5B). When we examined individual cells we found in 35% (n=46, 30 minutes) and 21% (n=28, 45 minutes) of cells Fir1 was present at the bud neck with no detectable Myo1.

**Figure 5:**
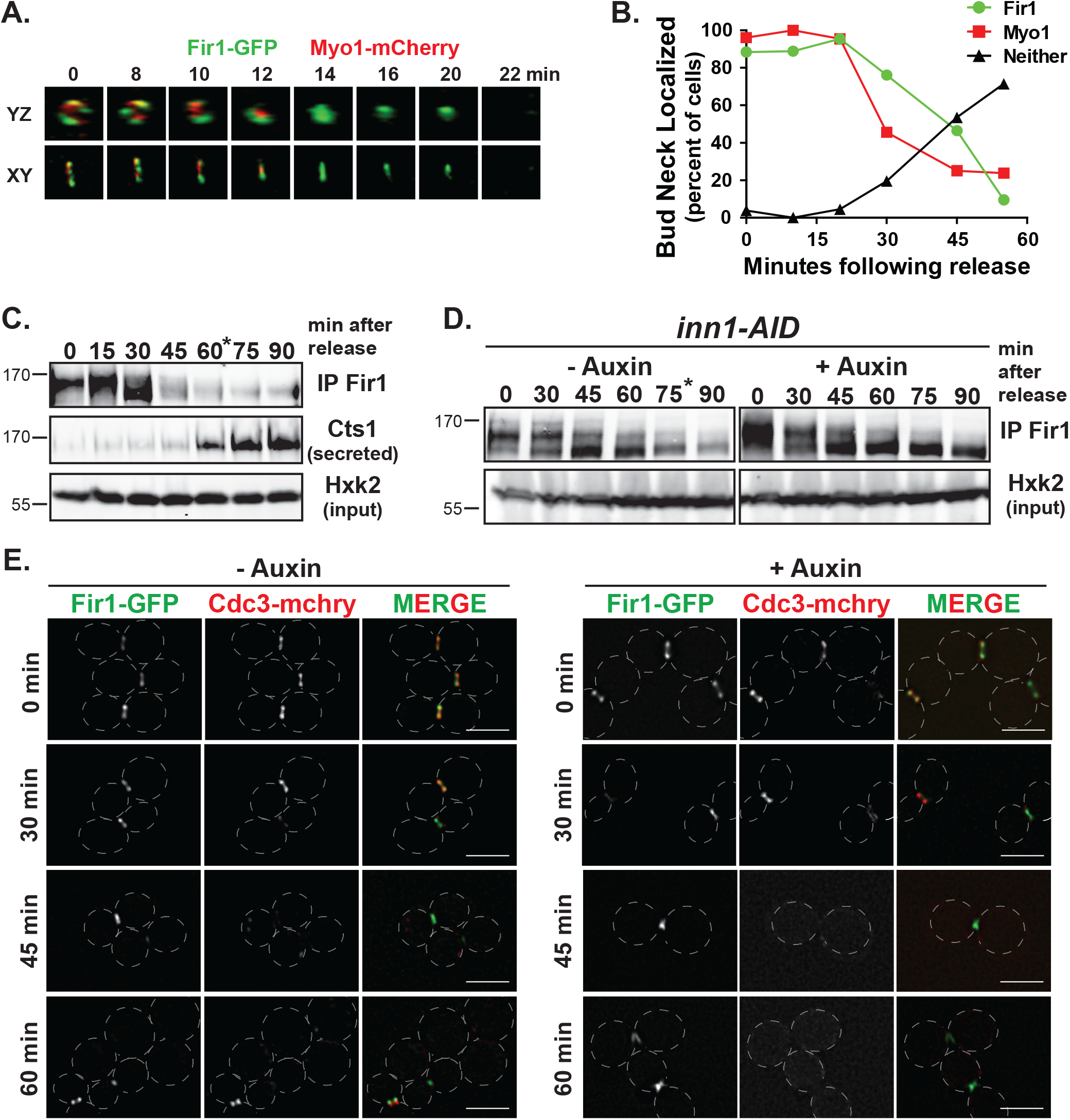
Fir1 is stabilized at the bud neck when septation is disrupted. (A) Fir1 localizes to the bud neck in late mitosis. Asynchronous cells expressing Fir1-GFP and Myo1-mCherry were subject to time lapse microscopy. Maximum projection of serial Z-stacks are shown in the XY and YZ planes at the times indicated after beginning image acquisition. A representative cell is shown. (B) Fir1 remains at the site of septation after the completion of cytokinesis. An aliquot of cells from a synchronized population were removed every 10 minutes after release and were imaged. The percent of cells with bud neck localization of the indicated protein is shown (n>20 cells per time point). (C) Fir1 is degraded prior to Cts1 secretion. Fir1-myc was immunoprecipitated (IP) from normalized lysates (input Hxk2 blot) following release from a synchronized culture at the times indicated at 30° C and subject to immunoblot. From the same culture, secreted Cts1 was collected and subject to immunoblot. A representative blot is shown. The asterisk indicates when peak cell separation was observed. (D) Fir1 is not degraded when septation is disrupted. Fir1-myc was immunoprecipitated from normalized lysates (Hxk2 input) in synchronized *inn1-AID* cells treated with or without auxin at 30° C and subject to immunoblot. The asterisk indicates when peak cell separation was observed. Since *inn1-AID* cells treated with auxin do not separate, no asterisk is shown. (See also Figure S4A). (E) Fir1 remains localized to the bud neck when septation is disrupted. Synchronized *inn1-AID* cells treated with or without auxin were released from arrest and representative images are shown from the indicated time after release. The septin mCherry-Cdc3 was used to mark the bud neck. The scale bar represents 5 μm. (See also Figure S4B-D).

Fir1’s disappearance from the bud neck could be due to its re-location or degradation. Notably, Fir1, unstable in G1, is a predicted target of the Anaphase Promoting Complex (APC) - Cdh1 (Ostapenko et al., 2012) and the Skp1-Cul1-F box (SCF) - Grr1 (Mark et al., 2014) ubiquitin ligases. To determine if Fir1 levels change as cells pass from late mitosis into G1, we synchronized cells by mitotic arrest and release and simultaneously examined Fir1 protein levels, cell separation, and Cts1 secretion. Fir1 exhibited significant electrophoretic shift at mitotic arrest, when Clb2-Cdc28 activity is high (metaphase / anaphase; T=0), and lost this shift as cells progressed through mitotic exit (Figure 5C). This presumably reflects phosphorylation and dephosphorylation, consistent with evidence that Fir1 is a target of the mitotic cyclin dependent kinase Clb2-Cdc28 (Holt et al., 2009; Kõivomägi et al., 2013) and the phosphatase Cdc14 (Kuilman et al., 2015). At 45 minutes after release from arrest, the amount of Fir1 present dropped significantly. This loss of Fir1 signal occurred just prior to peak cell separation (at 60 minutes) and importantly preceded Cts1 secretion (Figure 5C).

To assess the relationship of Fir1’s degradation to completion of septation, we examined Fir1 levels in *inn1-AID* cells treated with auxin and released from mitotic arrest. Unlike the mock-treated cells, Fir1 levels did not drop appreciably in auxin-treated cells (Figure 5D). In contrast, we found that the kinetics of Fir1’s loss of electrophoretic mobility shift were unaffected, suggesting efficient dephosphorylation (Figure 5D). We saw similar results in cells lacking the cytokinetic factor *CYK3* (Figure S4A), indicating that Fir1 stabilization is probably a consequence of cytokinetic failure and not specifically due to loss of Inn1.

We next examined Fir1-GFP localization in cells experiencing septation failure. All metaphase-arrested cells had Fir1-GFP at the bud neck. In the absence of auxin, the fraction of large-budded cells with bud neck Fir1-GFP dropped after release from arrest: 53% (n=80) at 45 minutes and 31% (n=91) of large budded cells 60 minutes after release. In contrast, 75% (n=96) and 76% (n=92) of large budded cells treated with auxin at 45 and 60 minutes retained Fir1 at the bud neck (Figure 5E). We also observed Fir1 localizing to the emerging bud in the next cell cycle; a localization pattern that was rarely seen in wild type or mock-treated cells (data not shown), consistent with reduced degradation upon failed cytokinesis. We saw a similar result in additional genetic backgrounds that are also defective in cytokinesis (Figure S4B and C). Taken together, these data suggest cells with defective septation activate a mechanism to stabilize Fir1 and retain it at the bud neck.

### Fir1 may inhibit the cell separation kinase Cbk1

The RAM (<u>R</u>egulation of <u>A</u>ce2 and <u>M</u>orphogenesis) signaling pathway initiates the cell separation process, and we hypothesized it might be subject to regulation by a septum monitoring system. The downstream most kinase, Cbk1, acts in a feed-forward loop to promote cell separation by regulating the transcription and translation of cell separation enzymes including *CTS1* (Jansen et al., 2009; Mazanka et al., 2008; Weiss, 2012). To test if elimination of *CBK1* would rescue the *inn1-AID* phenotype, we deleted *CBK1* in the *inn1-AID* strain. As expected, elimination of the kinase that initiates the cell separation process rescued *inn1-AID*’s growth defect (Figure 6A). To determine the epistatic relationship between *CBK1* and *FIR1*, we knocked out both genes and found this genotype suppressed the lethality of *inn1-AID* cells suggesting *CBK1* is epistatic to *FIR1* (Figure 6A). These data are likely due to lack of *CTS1* expression in *cbk1*Δ cells, however we cannot rule out the possibility that *FIR1* may act upstream of *CBK1* to block Cts1 secretion.

**Figure 6:**
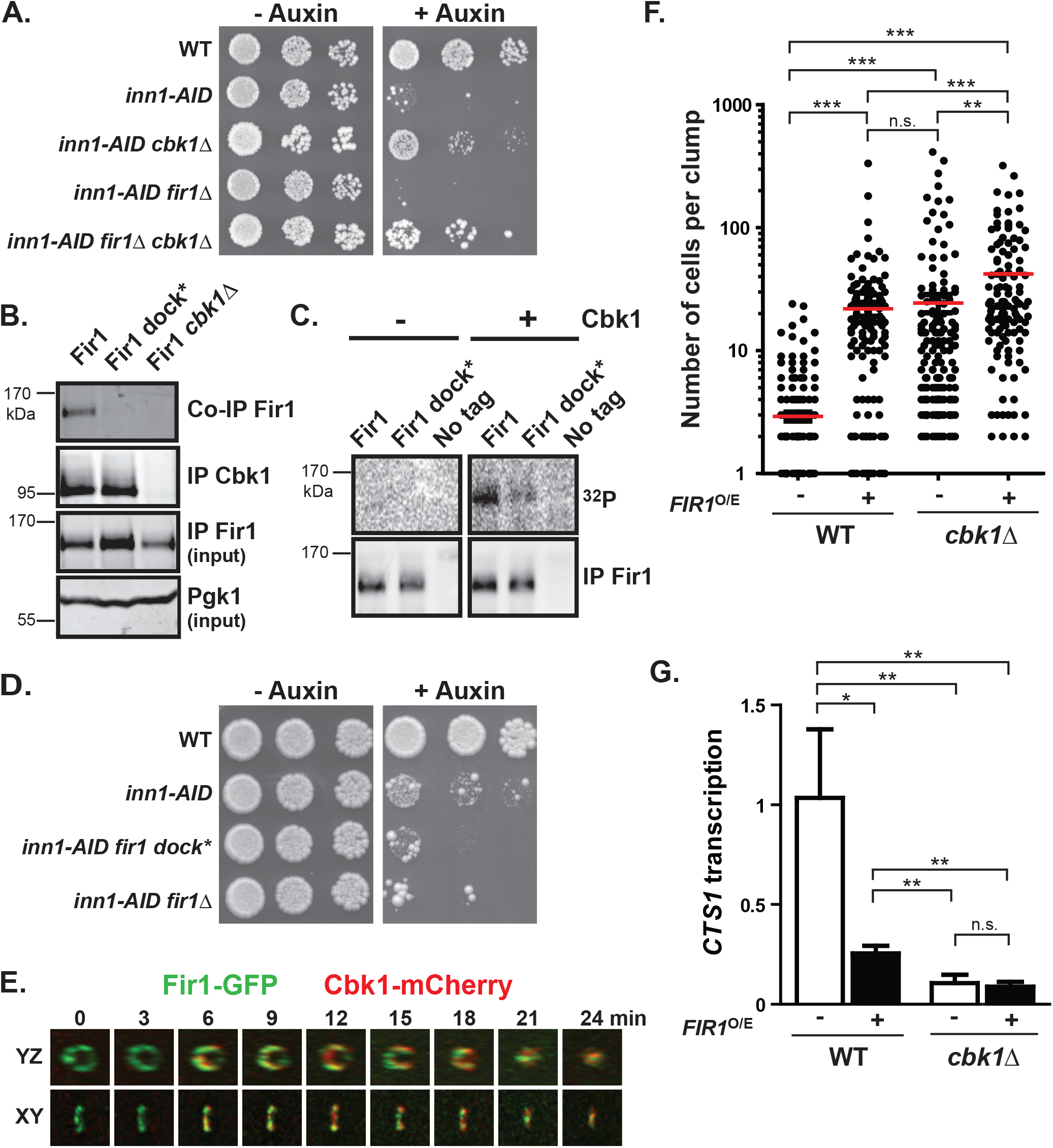
Fir1 may inhibit separation through inhibition of Cbk1. (A) *CBK1* deletion rescues the growth defect of cells that fail septation. Five-fold serial dilutions of the indicated strains were spotted to YPD media as in Figure 1C. (B) Fir1 interaction with Cbk1 *in vivo* requires Firl’s docking motif. Cbk1 was immunoprecipitated (IP Cbk1) from asynchronous cultures expressing wild type Fir1 or the Cbk1 docking motif mutant (Fir1 dock*). Immunoblot of the IP sample demonstrates Fir1 co-immunoprecipitation with Cbk1 (Co-IP Fir1). As a control, cells lacking *CBK1* were subject to the same IP and western analysis. Total Fir1 (IP Fir1 input) was examined by IP from the same normalized lysate (Pgk1 input). (C) Cbk1 phosphorylates Fir1 *in vitro*. HA-tagged Fir1, Fir1 dock* or an untagged strain were subject to Fir1 immunoprecipitation (IP) from asynchronous culture lysate and treated with or without bacterially purified Cbk1/Mob2 and radiolabeled ^32^P-ATP. An autoradiograph of Fir1 phosphorylation (upper panel) and an immunoblot showing total Fir1 (lower panel) is shown. (D) Fir1 function requires its interaction with Cbk1. Three-fold serial dilutions of the indicated strains were spotted to YPD media as in Figure 1C. (E) Cbk1 and Fir1 exhibit unique localization patterns during late mitosis. Synchronized cells expressing Fir1-GFP and Cbk1-3X mCherry were imaged every 3 minutes. Maximum projection of serial Z-stacks are shown in the XY and YZ planes at the time indicated after beginning image acquisition. A representative cell is shown. (See also Figure S5B). (F) *FIR1* overexpression inhibits cell separation. Wild type or *cbk1*Δ cells expressing an inducible *FIR1* overexpression vector (see methods) were grown in the absence or presence of inducer (−/+ *FIR1* O/E) at 30° C. The number of connected cells per cell clump (n>100 clumps) was counted for each sample. The red line indicates the mean clump size for the indicated strain. p-values are as follows: * = 0.01 to 0.05 ** = 0.01- 0.001 *** <0.001 (one-way ANOVA). (See also Figure S5C). (G) *FIR1* overexpression reduces Ace2 transcriptional output. RNA isolated from the cells in (F) were subject to qPCR analysis of *CTS1* transcript levels (normalized to *ACT1). CTS1* transcript levels are shown relative to *cbk1*Δ cells in the absence of inducer. Mean and standard deviation of three independent experiments are shown. p-values are as follows: * = 0.01 to 0.05; ** = 0.01- 0.001; n.s > 0.05 (t-test).

Recently, we identified a novel docking motif used by Cbk1 to interact with its substrates (Gógl et al., 2015). Fir1 contains two docking sites matching this consensus, [F/Y]XFP: one of which is highly conserved through fungal evolution (Nguyen Ba et al., 2012), Figure S2A i). Fir1 robustly co-immunoprecipitated with Cbk1 *in vivo* (Figure 6B, (Breitkreutz et al., 2010), and was not pulled down in the absence of *CBK1*. To test if this interaction required intact docking, we mutated the two [F/Y]XFP motifs to AXAP (Fir1 dock*). Fir1 abundance is low in the cell and we were unable to detected it in cell lysate. Therefore, we verified expression levels upon immunoprecipitation from normalized lysates (Figure 6B). While Fir1 dock* was expressed to a slightly higher level than wild type Fir1, elimination of the docking sites strongly reduced its interaction with Cbk1 (Figure 6B). Additionally, Fir1 contains several potential Cbk1 consensus phosphorylation sites (Figure S2, ii) that are also highly conserved through fungal evolution (Mazanka et al., 2008; Nguyen Ba et al., 2012). Fir1 immunoprecipitated from yeast lysate could be phosphorylated by bacterially expressed Cbk1 *in vitro*, and robust phosphorylation required an intact docking site (Figure 6C). Finally, to determine if Fir1’s function required its interaction with Cbk1, we examined the growth defect of *inn1-AID* when the endogenous *FIR1* allele was replaced with *fir1 dock\**. We found *fir1 dock\** did not restore full *FIR1* function to cells failing septation; cells grew more poorly in the presence of auxin exhibiting a phenotype more similar to *inn1-AID fir1*Δ cells (Figure 6D). Taken together, these data demonstrate Cbk1 and Fir1 are *bona fide* interaction partners and Fir1 function requires binding and/or phosphorylation by Cbk1.

Both Fir1 and Cbk1 localize to the site of septation and we wondered if altered localization could explain our genetic results. However, Fir1 and Cbk1 localized to the bud neck independently; both proteins were present at the bud neck in the absence of the other (Figure S5A). Then, we investigated their dynamics during cytokinesis. We found Fir1 forms a ring at the bud neck prior to Cbk1’s localization to the same ring structure (Figure 6E). Then, they fill in a disc-like structure as cells complete cytokinesis (Figure 6E, Movie S2, S3 and (Mancini Lombardi et al., 2013)). To clarify the timing of these proteins during cell separation, we examined localization in a synchronized cell population. At approximately the time of cell separation (50 minutes), a large proportion of cells (45%, n=56) exhibited bud neck localized Cbk1 in the absence of Fir1, suggesting Cbk1 remains at the bud neck after Fir1 degradation (Figure S5B).

Given the physical association between Fir1 and Cbk1, the timing of Fir1 and Cbk1 at the bud neck, and genetic evidence that Fir1 acts upstream of Cbk1 we investigated the possibility that Fir1 inhibits Cbk1 to prevent Cts1 secretion. Cbk1 inhibition prevents cell separation after septation as cells are unable to degrade the chitin-rich septum (Bidlingmaier et asdfal., 2001; Racki et al., 2000); therefore, we hypothesized the “clumpiness” or number of cells connected in a group would increase upon overexpression of an inhibitor. Wild type and *cbk1*Δ cells were transformed with an estradiol-inducible *FIR1* overexpression vector and treated with β-estradiol or vehicle (plus and minus Fir1 overexpression). Upon *FIR1* overexpression (see Figure S5C), cells exhibited a significant increase in the mean number of connected cells per group compared to cells without inducer, similar to cells lacking *CBK1* (Figure 6F). Since Cbk1 activates the transcription factor, Ace2, to drive transcription of *CTS1*, we also found *FIR1* overexpression reduced transcript levels of *CTS1* over 2-fold compared to mock treated cells (Figure 6G). Interestingly, we found *FIR1* overexpression in *cbk1*Δ cells lead to a slight but significant increase in the mean number of connected cells per group, suggesting Fir1 may have additional roles in preventing cell separation independent of Cbk1 (Figure 6F).

It was recently shown that the protein Lre1 can inhibit Cbk1 *in vitro*, and may also do so *in vivo* (Mancini Lombardi et al., 2013). We predicted that eliminating both putative inhibitors (Fir1 and Lre1) would additively increase Cbk1 activity, worsening viability defects of cells with impaired cytokinesis. Thus, we examined growth of *inn1-AID fir1*Δ and *inn1-AID fir1*Δ *lre1*Δ cells in the presence of auxin. We found no genetic interaction between *fir1*Δ and *lre1*Δ in the absence of auxin (no cytokinetic defect) (Figure S6A). At 30° C, both *inn1-AID fir1*Δ and *inn1-AID fir1*Δ *lre1*Δ strains grew extremely poorly, preventing meaningful assessment of genetic interaction (Figure S6A). However, we found that growth at 37° C partially restored growth defects of *inn1-AID* and *inn1-AID fir1*Δ cells (Figure S6A). In contrast, *inn1-AID fir1*Δ *lre1*Δ cells grew extremely poorly at 37° C, demonstrating a negative genetic interaction between *fir1*Δ and *lre1*Δ. Moreover, combining the *fir1 dock\** mutant with *inn1-AID lre1*Δ background demonstrated that the *fir1 dock\** allele was unable to restore growth to the *inn1-AID lre1*Δ cells at 37° C (Figure S6B), providing additional evidence that a physical association with Cbk1 is required for Fir1 function.

### Cbk1 regulates secretion of Cts1

We propose that Fir1 inhibition of Cbk1 helps prevent secretion of Cts1 before completion of septation. However, global inhibition of Cbk1 blocks Ace2-driven gene expression, and we found normal *CTS1* transcription in cells undergoing cytokinetic failure (Figure 2C). Thus, Fir1 may act at the bud neck to prevent Cts1 secretion through inhibition of Cbk1, and may not alter Cbk1’s ability to activate the transcription factor, Ace2. This model predicts that Cbk1 promotes Cts1 secretion more directly, in addition to its well established role in activating Ace2 to drive *CTS1* transcription (Mazanka et al., 2008; Racki et al., 2000; Weiss et al., 2002). To test this model, we examined Cts1 secretion in cells lacking *CBK1*. Since *CTS1* is not transcribed in *cbk1*Δ cells (Figure 6G and (Racki et al., 2000)), we replaced the endogenous *ACE2* with a gain-of-function *ACE2* allele (*ACE2-GOF*) that mimics Cbk1 phosphorylation by aspartic acid replacement at S122, S137 and S436 (Figure 7A and (Mazanka et al., 2008)). While this strain expresses *CTS1* to a lower level than wildtype *ACE2, CTS1* transcripts are produced with similar timing and levels in both *ACE2-GOF* and *ACE2-GOF cbk1*Δ (Figure S7A). We next assessed the production and secretion of Cts1 protein following release from mitosis in the presence and absence of Cbk1. We found that wild type and *cbk1*Δ cells carrying the ACE2-GOF allele had no difference in the production of intracellular Cts1 protein (Figure 7B and C). However, *cbk1*Δ cells had consistently reduced secretion of Cts1 secretion to the media (Figure 7B and C). This result suggests a novel Cbk1 function to promote Cts1 secretion independent of its role in promoting the transcription and translation of *CTS1*.

**Figure 7:**
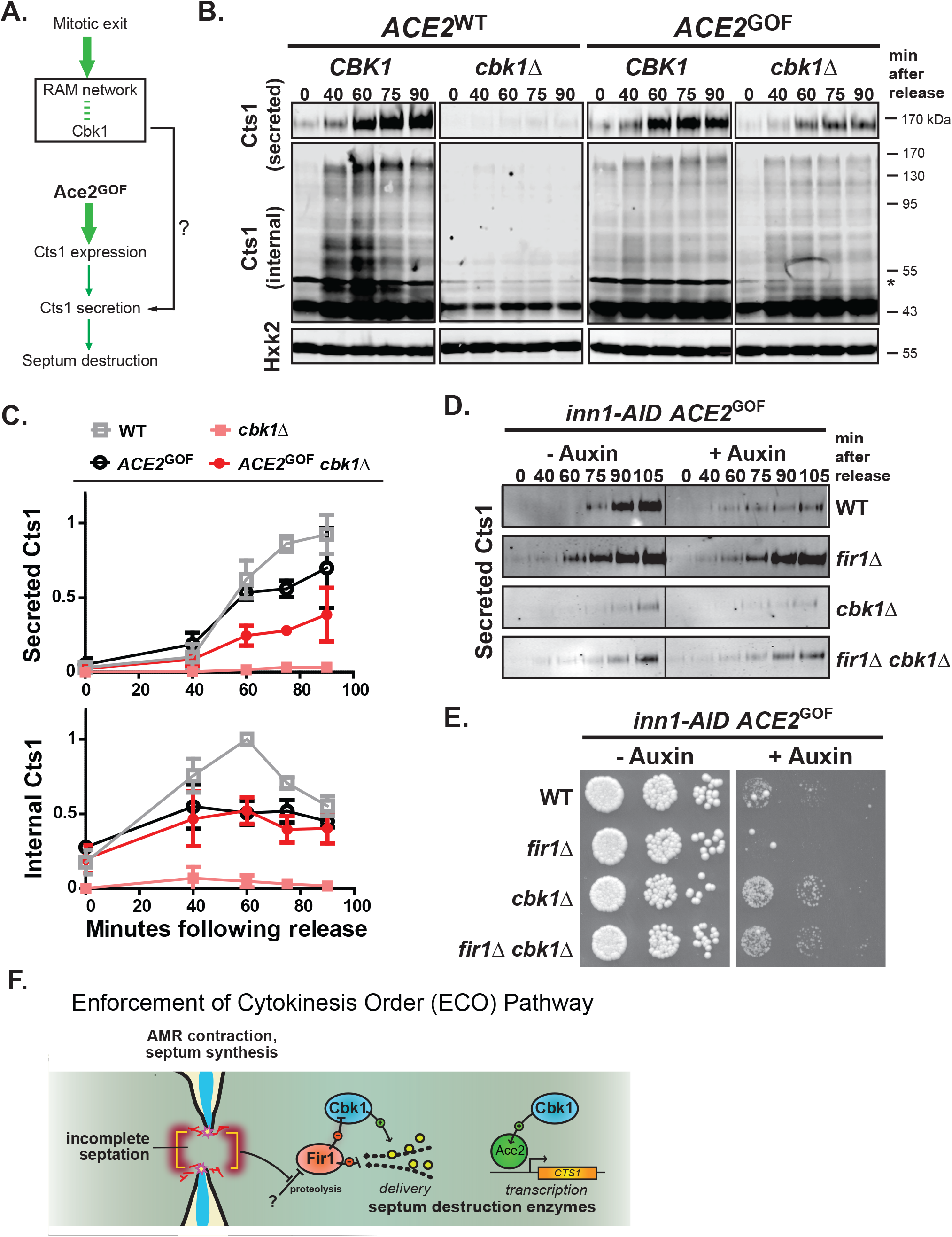
Cbk1 promotes Cts1 secretion independent of *CTS1* transcription. (A) Schematic of the *ACE2* gain-of-function (GOF) allele bypassing the necessity of Cbk1 to activate *CTS1* transcription. The *ACE2*-GOF allele mimics Cbk1 phosphorylation at S122, S137 and S436 to promote *CTS1* expression in the absence of *CBK1*. (B) Cts1 secretion is reduced in the absence of *CBK1* despite normal Cts1 production. Wild type, *cbk1*Δ, *ACE2*-GOF, *ACE2*-GOF *cbk1*Δ cells expressing HA-tagged Cts1 were synchronized in mitosis. Protein was collected at the indicated times following mitotic release at 30° C and were processed as in Figure 2A. (See also Figure S6A). (C) Western blot quantification (as in Figure 2B). (D) Fir1 primarily functions to inhibit Cbk1 to prevent Cts1 secretion. *inn1-AID ACE2*-GOF cells with the additional genotype indicated to the right of the panel were treated with DMSO (-auxin) or 0.5 mM auxin (+auxin). Secreted protein was collected at the indicated times following mitotic release at 30° C and a representative Western blot of secreted Cts1 is shown. (See also Figure S6B). (E) Cbk1 deletion restores viability upon septation failure despite Cts1 expression. Fivefold serial dilutions of the strains in (D) (all strains express PrGal-*CDC20 inn1-AID ACE2-GOF TIR1*) were spotted to YP Galactose media with and without the addition of 0.5 mM auxin. Plates were incubated for 2 days at 30° C. (F) “Enforcement of Cytokinesis Order” (ECO) pathway. Upon incomplete septation, stabilized Fir1 at the bud neck inhibits Cbk1, blocking secretion of septum destroying enzymes. Activation of ECO protects cytokinesis by ensuring the strict temporal sequence of opposing processes: septation and cell separation.

To validate this function of Cbk1, we also investigated the role of Cbk1 upon cytokinetic failure when *CTS1* is uncoupled from Cbk1 and expressed via the *ACE2-GOF* allele. Arrested *inn1-AID* cells expressing the *ACE2-GOF* allele were treated with or without auxin, released from the arrest, and Cts1 secretion was examined. Cts1 internal production is similar in all strains (Figure S7B). As shown previously, cells failing septation do not secrete Cts1 (Figure 7D, WT) while cells lacking *FIR1* bypass the block and secrete Cts1 (Figure 7D, *fir1*Δ). Importantly, cells lacking *CBK1* secrete very little Cts1 regardless of cytokinesis failure (Figure 7D, *cbk1*Δ, +/− auxin). These data demonstrate functional Cbk1 is required for Cts1 secretion and is consistent with our model that Cbk1 inhibition is important for prevention of Cts1 secretion. Moreover, we find that additionally eliminating *FIR1* only slightly increased the amount of Cts1 secreted to the media (Figure 7D, *fir1*Δ *cbk1*Δ). These data uphold our genetic evidence that *CBK1* is epistatic to *FIR1* and provide additional evidence that Fir1 has only a minor role in preventing Cts1 secretion independent from its inhibition of Cbk1. Consistent with these results, we found *CBK1* deletion rescued the growth defect of *inn1-AID* and *inn1-AID fir1*Δ cells that express *CTS1* independent of *CBK1* (Figure 7E). Taken together, these results demonstrate a novel Cbk1 function to promote Cts1 secretion and suggests Fir1 inhibition of Cbk1 could prevent Cts1 secretion when septation is disrupted.

## Discussion

Our findings indicate that the final stages of budding yeast cytokinesis are sensitive to the status of the preceding stages of the process. This is manifested by pronounced delay in secretion of proteins that destroy the septum when septum completion is delayed or disrupted. We propose that a checkpoint-like “Enforcement of Cytokinesis Order” (ECO) pathway (Figure 7F) maintains the strict temporal sequence of the incompatible stages of cytokinesis, septation and cell separation. Dependency of septum destruction on completion of cytokinesis requires the protein Fir1, which inhibits secretion of the septum degradation enzyme Cts1: notably, Fir1 is not essential during unperturbed division. Septum completion and abscission appear to trigger destruction of Fir1, relieving the block to septum destruction. Our evidence suggests the ECO pathway blocks premature Cts1 secretion in two distinct ways: by directly inhibiting Cts1 secretion, and by inhibiting the protein kinase Cbk1, the Ndr/Lats component of the conserved RAM network ‘hippo’ pathway. We find that Cbk1 is crucial for normal Cts1 secretion septum, a previously unappreciated function that is separate from its well-defined role in activation of CTS1 transcription. We propose that the ECO pathway functions by stabilizing Fir1 in cells that have not executed septation and abscission, and that completion of these stages of cytokinesis triggers destruction of Fir1 is to allow secretion of septum-degrading proteins.

### A checkpoint that monitors cytoplasmic separation

Canonically, checkpoints are defined as systems that create and enforce dependency relationships for cell cycle processes in which late events would otherwise occur without regard to completion of earlier ones (Hartwell and Weinert, 1989). The ECO pathway appears to protect budding yeast cytokinesis by ensuring that the onset of cell separation does not occur before septation is complete. In some cases, a checkpoint’s action is only strongly evident when the processes it monitors are defective. For example, components of the budding yeast DNA damage and spindle assembly checkpoints are nonessential under ideal growth conditions (Li and Murray, 1991; Weinert and Hartwell, 1988). Cells lacking Fir1 have no pronounced phenotype but are highly sensitive to treatments that disrupt cytokinesis. For example, cells experiencing conditional disruption of septation (*inn1-AID* cells treated with auxin) quickly lose viability in the absence of Fir1, failing to delay Cts1 secretion and exhibiting phenotypes consistent with failed cytokinesis and premature septum destruction.

Our findings are consistent with prior observations indicating that septum destruction depends on successful execution of cytokinetic abscission and septum completion: specifically, cells that undergo remedial septum formation do not perform mother-daughter separation (Atkins et al., 2013; Bi et al., 1998; Cabib, 2004; Onishi et al., 2013; Schmidt et al., 2002; Shaw et al., 1991; Yeong, 2005). This research broadly hypothesized that the relatively disorganized remedial septum is refractory to separation. However, we find that the remedial septum is destroyed upon ECO bypass. The mechanisms that link ECO to the mechanical progress of cytokinesis are unclear. Intriguingly, however, cells lacking Myo1 exhibit enhanced Cts1 secretion (Ríos Muñoz et al., 2003). We found that Fir1 and Myo1 closely colocalize in early cytokinesis, and speculate that cells lacking Myo1 fail to activate ECO. Of note, *myo1*Δ cells readily accumulate suppressive aneuploidies (Rancati et al., 2008; Tolliday et al., 2003), consistent with a detrimental effect of premature septum destruction, worsened by loss of mechanisms enforcing the dependency of separation on septation.

Like other checkpoints, the ECO pathway appears to protect cytokinesis by inhibiting a late event (septum destruction) until an earlier one (septation) is completed. The concept of early factors preventing the action of late factors has been demonstrated recently for the early events of cytokinesis. Recent work from several laboratories have described elegant mechanisms to ensure order during AMR contraction, septum formation and bud emergence (Atkins et al., 2013; Meitinger et al., 2013; Oh et al., 2017; Onishi et al., 2013). For example, Cyk3, a protein required for PS formation (early), was demonstrated to inhibit the Rho1 GTPase required for SS formation (late) (Onishi et al., 2013). However, we propose here a different mechanism to ensure order. Unlike these other factors, Fir1 does not appear to be required for any step of septation or cell separation. Cells lacking *FIR1* have no obvious phenotype. Instead, Fir1, like other checkpoint proteins, functions to relay information about the status of an early event to a late event. Upon septation failure, Fir1 is stabilized and transmits this information to the RAM network preventing later processes.

What is the signal that relays to Fir1? Checkpoints function by monitoring a process and sending a negative signal to block subsequent processes until a particular condition is satisfied. In ECO, we do not yet understand which aspect of septation is being monitored. Cells may monitor defects as cell wall stress via activation of the cell wall integrity pathway (Philip and Levin, 2001) or they may monitor disruptions to the plasma membrane (Kono et al., 2012) or they may monitor a yet unidentified morphological change. Interestingly, decreasing lipid flippase function at the plasma membrane during cytokinesis was shown to suppress the poor growth of septation mutants (Roelants et al., 2015) suggesting that plasma membrane composition changes during cytokinesis could be involved.

Intriguingly, the crucial ECO pathway component Fir1 is an “intrinsically disordered” protein (IDP) with no predicted folded domains but numerous conserved peptide motifs that are known or likely binding partners of folded protein domains (also known as short linear motifs or “SLiMs”) (Nguyen Ba et al., 2012). It is becoming clear that intrinsically disordered proteins (IDP) perform diverse important roles, in some cases by generating distinct “phase separated” regions (Woodruff et al., 2017; Wright and Dyson, 2014). Fir1 has an unusually large number of SLiMs (Nguyen Ba et al., 2012), suggesting that it may function as a signaling hub by concentrating multiple interaction sites (Dunker et al., 2005). Intriguingly, rapidly-evolving IDPs containing conserved SLiMs and small folded motifs are important in other conserved checkpoints and signaling systems. For example, the Rad9 protein - among the first components identified of the DNA damage checkpoint - is nearly entirely IDP, with a concentration of short BRCT motifs and SLiMs (Weinert and Hartwell, 1988).

### How does the ECO pathway protect cytokinesis?

Septation and cell separation in yeast are controlled by two deeply conserved ‘hippo’ pathway signaling systems: the MEN and the RAM networks (Bardin and Amon, 2001; McCollum and Gould, 2001; Weiss, 2012). Response to cytokinesis disruption probably does not involve the MEN, which acts prior to AMR contraction and septum initiation. We saw no delay of cytokinesis initiation in synchrony experiments, which is inconsistent with MEN inhibition. The RAM network’s best-known function in cell separation is activation of Ace2-driven transcription of cell separation genes. Delay of separation could therefore occur by suppression of Ace2 function through RAM network inhibition, as well as through block of cell separation gene translation or delivery of these enzymes to the nascent, unfinished septum. We find that both transcription and translation of the Ace2-driven gene *CTS1* is not affected when cytokinesis is impaired. Rather, cells prevent premature secretion of this enzyme, indicating that the membrane trafficking system is responsive to the status of cytokinesis.

Intriguingly, the ECO pathway may work in part by inhibiting RAM network functions that are independent of Ace2 activation. By unlinking Ace2 function from the RAM network we were able to determine that secretion of Cts1 is slowed when the RAM network is inhibited, suggesting that RAM plays a role in control of secretion during late cytokinesis. While this could reflect a total block in secretion during cytokinesis, we think this is unlikely. For one, remedial septum synthesis occurring at this time depends on secretion of Chs3 (Cabib and Schmidt, 2003), and we have also found the exocyst component, Sec4, still relocalized to the bud neck at this time (data not shown). Furthermore, it is unlikely that the RAM network plays a major role in secretion outside of cytokinesis. In addition to the brief event of cell separation, the RAM network plays substantial roles in cell growth and morphogenesis in other times during in the cell cycle. Specifically, cells lacking the pathway bud and proliferate with normal kinetics but do not sustain polarized growth (Bidlingmaier et al., 2001; Weiss et al., 2002). A significant part of this role is inhibition of translation repression of proteins that are required for wall expansion (Jansen et al., 2009; Wanless et al., 2014). In cells lacking this translation control system, disruption of RAM network function affects maintenance of polarized growth but has no pronounced effect on proliferation or bud growth. We, therefore, propose that the RAM network either promotes default secretion only during septation and cell separation, or specifically activates the trafficking/secretion of specific cargo proteins involved in cell separation. While Cbk1 may regulate glycosylation (Kurischko et al., 2008) we did not find obvious changes in Cts1 glycosylation in cells lacking *CBK1* (Figure 7B). Interestingly, the predicted Cbk1 target Boi1 (Gógl et al., 2015) has recently been shown to promote activation of the exocyst (Kustermann et al., 2017; Masgrau et al., 2017). While any requirement for the exocyst in promoting Cts1 secretion is unknown, secretion of septum destroying enzymes in fission yeast requires functional exocyst (Martín-Cuadrado et al., 2005). Overall, the broader significance of our findings is that trafficking of cytokinetic proteins is precisely timed according to their order of function.

Fir1 is clearly a target of Cbk1 that interacts with the kinase through a conserved docking motif, but the mechanism of inhibition by Fir1 remains unclear. Fir1 lacking functional docking sites did not restore viability to cells failing septation (*inn1-AID* treated with auxin) suggesting inhibition requires this interaction. One possibility is that Fir1 may physically block Cbk1’s interaction with other docking-motif containing targets until Fir1 degradation. We have computationally identified potential Cbk1 targets with strong conservation of docking and/or phosphorylation sites (Gógl et al., 2015). Interestingly, while most exhibit conservation of both docking and phosphorylation sites, several lack a conserved docking site. An intriguing possibility is that Fir1 can order Cbk1 substrates by permitting access to some substrates and not others during late stages of cytokinesis. However, we cannot rule out the equally likely, and not mutually exclusive, possibility that Cbk1 phosphorylation of Fir1 plays a more important role in its function. Perhaps phosphorylation of these sites enhances or decreases Fir1’s ability to inhibit Cbk1, and this is a key area of further investigation.

### How broadly conserved is the ECO pathway?

We suggest that checkpoint-mediated enforcement of dependency in cytokinesis similar to the ECO pathway in overall structure, if not in specific components, is broadly distributed in eukaryotes. Our results are notably consistent with experiments performed in fission yeast: uncoupling the RAM-related pathway, MOR (morphogenesis Orb6 network), from upstream controls caused cell lysis. The authors demonstrated that cells failing to inactivate the MOR early during septation prematurely degraded the septum leading to lethality. Similar to our results here, this phenotype was reversed by deletion of wall degrading enzymes (Gupta et al., 2014). While fission yeast lack a sequence homolog of Fir1, a functional ortholog may exist. We predict, however, this functional ortholog should prevent premature activation of the MOR pathway upon failed septation independent of upstream controls.

Fundamental aspects of cytokinesis are conserved between yeast and animal cells (Balasubramanian et al., 2004; Bhavsar-Jog and Bi, 2017; Glotzer, 2016). Both yeast and metazoan cells undergo cytokinesis through a sequence of coordinated steps that include AMR contraction, membrane ingression, localized secretion, extracellular matrix (ECM) remodeling, and abscission. Mechanisms to ensure coordinated execution of these steps are beginning to be understood (Glotzer, 2016; Neto and Gould, 2011). For example, kinases play a fundamental role in ensuring abscission order (Paolo D’Avino and Capalbo, 2016). The polo-like kinase Plk1, activate in late mitosis, prevents the Cep55 protein from prematurely localizing to the midbody. Upon mitotic exit, Plk1 activity drops and recruitment of Cep55 ensures timely assembly of the machinery required for abscission (Bastos and Barr, 2010). Then Cep55 recruits exocyst components to promote secretion required for abscission (Gromley et al., 2005; Zhao et al., 2006). Remodeling of the extracellular matrix is also important for proper cytokinesis (Hwang et al., 2003; Olson et al., 2006; Xu and Vogel, 2011). However, the dependency relationships between these processes: abscission, localized secretion and ECM remodeling is not known. Given the highly conserved nature of many other aspects of cytokinesis, it is interesting to speculate that, like the budding yeast ECO pathway, metazoan cells use similar checkpoint mechanisms to ensure order of these diverse processes.

## Methods

### Strains, plasmids, and growth conditions

All strains are derived from the W303 genetic background (*leu2-3,112 trp1-1 can1-100 ura3-1 ade2-1 his3-11,15*) and are listed in Supplemental Table 1. We generated deletion and C-terminal hemagglutinin (HA), c-Myc (Myc), green fluorescent protein (GFP), and red fluorescent protein (mCherry)-tagged strains by standard integration methods (Longtine et al., 1998; Sheff and Thorn, 2004). Integration of a 3X-HA internal tag into the Cts1 coding sequence was performed with a pop-in/pop-out method (Schneider et al., 1995) just after the catalytic domain of Cts1 between amino acids A315 and T316. We sequenced the endogenous locus to confirm proper integration. The dual (N- and C-terminally) tagged Fir1 and Fir1 dock* strains were generated by a two-fragment PCR method to replace the *fir1*Δ::KanMX with a PCR amplification of GFP or HA-Fir1 from a plasmid and Fir1-myc :: TRP1 from genomic DNA. Untagged *fir1 dock*\* was integrated at the endogenous locus replacing a CORE cassette using the Delitto Perfetto method (Storici and Resnick, 2006). We sequenced the locus to ensure proper integration. The ACE2-GOF (S122D, S137D and T436D) allele was integrated using a two-fragment PCR method to replace *ace2*Δ::*HIS3* with the gain-of-function allele, a GFP tag and the KanMX marker. The endogenous locus was sequenced to confirm proper integration. The septin Cdc3 was N-terminally tagged upon BglII digest of YIp128-CDC3-mCherry (*LEU2*, a gift from E. Bi (Gao et al., 2007)) and transformation. The plasma membrane marker PH-GFP (LEU2, a gift from Y. Barral (Mendoza et al., 2009)) was transformed into the indicated strains. Derivatives of the *inn1-AID PrADH1-O.s.TIR1-9MYC* strains were generated by backcrossing to strain YAD236 (a gift from K. Labib (Devrekanli et al., 2012)). Genotypes were determined by marker segregation or via PCR confirmation of unmarked alleles (Cts1 internal HA screened with primers: 5’ATTTGCTAACAAGTGCTAGCCAGAC 3’ and 5’ GATGAAGTTGAGGCTGCTGAGG 3’).

We generated *CTS1* and *FIR1* overexpression constructs using the Drag and Drop method (Jansen et al., 2005). Briefly, the coding sequence of *CTS1* or *FIR1* were PCR amplified with Rec1 and Rec2 overhangs and co-transformed with SalI digested pGREG505::*LEU2* or pGREG575::LEU2 vectors, respectively. Inserts were integrated into the vector by homologous recombination gap repair. The resulting plasmids were recovered and sequence confirmed (pELW 2038 and pELW 2039, respectively). The uncut vector served as an empty vector control.

We cultured cells in YP medium (1% yeast extract, 2% peptone; BD) containing additional adenine (Ameresco) and 2% glucose (EMD) (YPD) or 2% galactose (MP Biomedical) (YP Gal). For microscopy we used synthetic minimal selection medium (0.67% yeast nitrogen base without amino acids, 0.2% amino acid drop-in; US Biological, and 2% glucose or 2% galactose). Plated media was as above with 2% agar (US Biological). DMSO (Sigma) and auxin (α-naphthalene acetic acid, HiMedia Laboratories) were added to plates or media at 0.5 mM final concentration where necessary.

### Cell synchrony

For mitotic arrest and release, we grew cells expressing a galactose-inducible GalL-*CDC20* gene to log phase in YP Gal media, washed once with YP media, resuspended in fresh YPD to turn off expression of Cdc20 and arrested in metaphase. We incubated cultures at 30° C until > 95% of the cells were large budded (2.5 – 3 hours). For auxin depletion experiments, cultures were split equally and auxin (or DMSO) was added to 0.5 mM final concentration for the last 30-45 minutes of the arrest. We washed cells twice in YP media (containing auxin or DMSO where necessary) and released in YP Gal media (containing auxin or DMSO where necessary) at the temperature indicated in the figure legend. At each time point, a sample was removed and the budding index of 100 cells was counted to confirm arrest and release.

In Figure 4A and 4C, strains were engineered to induce GalL-*CDC20* with deoxycorticosterone (DOC) driven activation of Gal4 (Picard, 2000). We grew cells to log phase in the presence of 50 nM DOC (dissolved in ethanol), washed three times to remove hormone, and induced the metaphase arrest in the absence of hormine. We incubated cultures at 30° C until > 95% of the cells were large budded (2.5 – 3 hours). Cultures were then split equally and auxin (or DMSO) was added to 0.5 mM final concentration for the last 30 minutes of the arrest. To enrich for cells completing cytokinesis, we removed an aliquot of arrested cells and added 50 nM DOC to induce release. After an additional 10 and 20 minutes, another equal aliquot of cells from the arrest were added to this with additional DOC (to maintain 50 nM DOC). After 80 minutes, we fixed the mix of cells so that the mixture contained an equal proportion of cells released at 60, 70 and 80 minutes following addition of DOC. We then processed them for chitin staining and electron microscopy. Since minus auxin cells complete cytokinesis more rapidly, we collected cells as above but fixed cells after 60 minutes so that the mixture contained an equal proportion of cells released at 40, 50 and 60 minutes following addition of DOC.

### Preparation of secreted and internal Cts1 protein

To examine Cts1 protein production, we synchronized cells as indicated above. At the time points indicated in the figure, we removed an equivalent number of cells from each culture (typically 2 OD cell equivalents), added sodium azide (Sigma) (to 20 mM final concentration) and placed on ice to prevent further protein production or secretion. At the end of the time course, cells were spun down at 20,800 × g for 1 minute and 90% of the cleared media was added to ice-cold trichloroacetic acid (TCA, Sigma) (10% final concentration) and we precipitated secreted protein overnight at 4° C. Precipitated protein was pelleted at 20,800 × g for 15 minutes at 4° C, washed once in ice-cold acetone (Sigma) and pellets dried at room temperature. Samples were resuspended in 2X buffered protein loading buffer (125 mM Tris pH 9.4, 20% glycerol, 4% SDS, 10% 2-mercaptoethanol, 0.004% bromophenol blue) and boiled at 99° C for 5 minutes. We ran half of the sample on a 10% SDS-PAGE gel and immunoblotted the gels as indicated below.

We washed the remaining cells into 1.2 M sorbitol, 10mM Tris pH 7.6 buffer, then incubated with 0.5 mg/ml Zymolyase 20T (ICN) in 1.2 M sorbitol, 10mM Tris pH 7.6, 50 mM 2-mercaptoethanol for 30-40 minutes at 37° C to remove the cell wall. Spheroplasts were gently centrifuged at 830 × g for 1 minute. Cells were lysed on ice by resuspension and brief vortex in 50 mM Tris pH 7.5, 0.5mM EDTA, 0.5% Triton X-100, 1 mM PMSF. We removed cell debris upon centrifugation at 20,800 × g for 15 minutes at 4° C. The cleared lysate was added to 2X protein loading buffer (125 mM Tris pH 6.8, 20% glycerol, 4% SDS, 10% 2-mercaptoethanol, 0.004% bromophenol blue) and boiled at 99° C for 5 minutes. Equal volumes (10-20% of the total lysate) were loaded on a 10% SDS-PAGE gel and subject to immunoblotting as indicated below.

### Immunoprecipitation

We prepared cell lysate by bead beating in ice-cold lysis buffer (50 mM Tris-HCl pH 7.4, 150 mM NaCl, 1% Triton X-100, 10% glycerol, 1 mM dithiothreitol, 120 mM β-glycerolphosphate, 2 mM sodium orthovanadate, 20 mM sodium molybdate, 3 mM benzamidine, 1 mM phenylmehtylsulfonyl fluoride, 1 ug/ml pepstatin, 0.5 mM leupeptin, and 2 ug/ml chymostatin). Anti-HA antibody (12CA5, a gift from R. Lamb) at a 1:50 dilution along with 50 ul of a 1:1 recombinant protein G-Sepharose (Invitrogen) slurry was added to normalized protein, as determined by Bradford assay (Bio-Rad). We rotated immunoprecipitations at 4°C for 2 h followed by two washes in ice-cold yeast lysis buffer. We then resuspended the beads in 2x SDS-PAGE sample buffer (125 mM Tris pH 6.8, 20% glycerol, 4% SDS, 10% 2-mercaptoethanol, 0.004% bromophenol blue) and resolved the samples on 8% SDS-PAGE gels. Gels were immunblotted as indicated below.

### Co-immunoprecipitation

We prepared cell lysates of asynchronous cells by bead beating in ice-cold lysis buffer (50 mM Tris-HCl pH 7.4, 150 mM NaCl, 1% Triton X-100, 10% glycerol, 1 mM dithiothreitol, 120 mM β-glycerolphosphate, 2 mM sodium orthovanadate, 20 mM sodium molybdate, 3 mM benzamidine, 1 mM phenylmehtylsulfonyl fluoride, 1 ug/ml pepstatin, 0.5 mM leupeptin, and 2 ug/ml chymostatin). We normalized total protein as determine by Bradford assay (Bio-Rad) and split the lysate equally into two. A sample was removed and added to sample buffer for the input loading control. One aliquot was incubated with anti-HA antibody (12CA5, a gift from R. Lamb) and recombinant protein G-Sepharose (Invitrogen) slurry to determine the total Fir1-HA in the lysate. The other aliquot was incubated with anti-Cbk1 NT5 antibody (Jansen et al., 2006) and recombinant protein G-Sepharose (Invitrogen) slurry for the co-immunoprecipitation. We rotated the beads at 4°C for 2 h followed by three washes in ice-cold yeast lysis buffer. We then resuspended the beads in 2x SDS-PAGE sample buffer (125 mM Tris pH 6.8, 20% glycerol, 4% SDS, 10% 2-mercaptoethanol, 0.004% bromophenol blue) and resolved the samples on 8% SDS-PAGE gels. Gels were immunoblotted as described below.

### Immunoblotting

For all Western blot assays, we transferred proteins to Immobilon FL PVDF (polyvinylidene difluoride; Millipore) membrane. Membranes were blocked for 30 minutes in Odyssey blocking buffer PBS (Licor), then incubated with primary antibody in Tris-buffered saline (50 mM Tris pH 7.6, 150 mM NaCl) plus 0.1% Tween (TBST) for 1 h at room temperature or overnight at 4° C. We then incubated blots with secondary antibody in TBST plus 0.1% SDS for 30 minutes. Membranes were washed three times for 3 minutes with TBST between antibody additions and prior to imaging. We used primary antibodies as follows: mouse monoclonal HA antibody (12CA5; a gift from R. Lamb) at 1:1000, rabbit polyclonal hexokinase (Rockland Immunochemicals) at 1:3000, and rabbit polyclonal Clb2 antibody (y-180, Santa Cruz Biotech) at 1:1000. We used secondary antibodies as follows: IRDye 680 LT goat anti-mouse (Licor) at 1:20,000 or IRDye 800 CW goat anti-rabbit (Licor) at 1:15,000. We imaged and quantified blots with the Image Studio Lite Odyssey software (v4.0; Li-Cor), and images of blots were processed using Photoshop (Adobe). Quantification was normalized by scaling intensity data from 0 to 1 using the equation x’ = (x-min(x)) / (max(x) – min (x)). The min and max values were determined from each independent time course experiment and included all data from inn1-AID (Figure 2) and inn1-AID fir1D (Figure 3) or from wild type, *cbk1*Δ, *ACE2-GOF, ACE2*-GOF *cbk1*Δ in Figure 7. Internal Cts1 was additionally normalized by generating a normalization factor from the Hxk2 blot (x/max(x)) and dividing the raw Cts1 value by this before scaling.

### Kinase assay

We performed kinase assays as previously described (Jansen et al., 2006). Briefly, we immunoprecipitated Fir1-HA from yeast lysate (see above) and washed the beads three times with kinase reaction buffer (20 mM Tris, pH 8.0, 150 mM NaCl, and 5 mM MnCl2, 20 mM DTT). We resuspended the beads in reaction buffer plus 20 μM cold ATP and 0.33 μCi/μL γ-^32^P-ATP and added 240 nM bacterially purified Cbk1^251–756^ - Mob2 (Gógl et al., 2015). Kinase reactions were allowed to proceed at room temperature for 1 hour and stopped by addition of 5× SDS-PAGE sample buffer and 20 min incubation at 80°C. Proteins were separated by 8% SDS-PAGE and transferred to Immobilon FL PVDF (polyvinylidene difluoride; Millipore) membrane. We visualized ^32^P using a Storm phosphorimager (GE Biosciences) and total Fir1 was imaged by immunoblotting as described below.

### RNA preparation and QPCR

We prepared RNA from synchronized cultures at the times indicated in the figure by hot acid phenol extraction as previously described (Collart et al.). We treated 2 μg of RNA with 10 units of RNase-free DNase I (Roche) and converted it to cDNA with Moloney murine leukemia virus reverse transcriptase (MMLV-RT; Promega). We performed quantitative RT-PCRs (qPCR) with the iCycler Thermal Cycler with iQ5 Multicolor Real-Time PCR Detection System (Bio Rad) using primers specific to *CTS1* and *ACT1* (*CTS1* Fwd: 5’ TGCACCCAGATTGCTGAA 3’ Rev: 5’ AAACCATCAACGACTGCTGAG 3’ and *ACT1* Fwd: 5’ GGTTATTGATAACGGTTCTGGTATG 3’ Rev: 5’ ATGATACCTTGGTGTCTTGGTCTAC 3’). Efficiencies (E) were obtained by plotting the log of the relative template concentration to the cycle threshold (CT) values from serial dilutions of yeast genomic DNA and calculating 10^−1/(slope-1)^. Efficiency corrected gene expression levels were calculated using the ratio of *CTS1* over *ACT1* with the equation: E^ΔCT target (T=0 – T= × min sample)^ / E^ΔCT ACT1 (T=0 – T= × min sample)^. Fold changes are shown relative to the control sample indicated in the figure legend. A student’s unpaired, twotailed t-test was performed using GraphPad Prism version 5.03 to determine significance.

### Spot assay and cell separation assays

For spot assays, we grew cells in YPD to mid-log phase then normalized to an OD 600 of 0.05 in water. Five-fold or three-fold (see figure legend) serial dilutions were made and three microliters were spotted onto the plates. We incubated plates for 3-5 days at 30C (unless otherwise indicated). For cell separation assays, wild type or *cbk1*Δ cells were transformed to integrate PYL23 (PrADH1 Gal4 ER VP16) at the *URA3* locus upon digestion with Xcm1 (gift from F. Cross). This generates a strain in which the Gal4 transcription factor is activated upon addition of β-estradiol (Picard, 2000). These strains were then transformed to express FIR1 from the Gal1 promoter (pELW 2039, see above). Finally, the resulting strains were grown in selection media to mid-log phase and diluted again to allow growth to log-phase overnight (16 hours) in the absence or presence 125 nM β-estradiol (−/+ *FIR1* O/E). Samples were randomized, imaged and analyzed blind. We imaged more than 100 cell clumps per sample and the number of cells in each clump were plotted. The mean cell per clump is indicated as a red line in the figure. One-way ANOVA with Tukey’s Multiple Comparison Test was performed using GraphPad Prism version 5.03.

### Chitinase enzymatic assay

We washed metaphase arrested cells three times in fresh media to eliminate previously secreted Cts1. Cells were released into fresh galactose containing media and grown at 30° C. At the times indicated in the figure, we treated an aliquot of cells with sodium azide (to 10mM final concentration) and placed on ice to prevent further secretion. We mixed 30 ul of cells and media with 20 ul of chitinase substrate (250 μM 4-Methylumbelliferyl β-D-N,N’,N”-triacetylchitotrioside (Sigma) in 0.25 M sodium citrate pH 3.0) and incubated the reaction at 30° C for 0, 30, 60 and 90 minutes. At these times, the reaction was terminated by addition of 50 ul 0.5 M glycine-NaOH, pH 10.5. Liberated 4-methylumbelliferone (4-MU) was measured with a Synergy 4 microplate reader (BioTek) at an excitation wavelength of 360 nm and an emission wavelength of 450 nm. We used a standard concentration of 4-methylumbelliferone (4-MU) to calculate the activity of chitinase (pmol 4-MU releasedl/min/cell OD) at each time point after metaphase release.

### Calcofluor stain

We fixed the indicated strain with 3.7% formaldehyde and washed twice in PBS (4mM KH_2_PO_4_, 16 mM Na_2_HPO_4_, 115 mM NaCl, pH 7.3). We resuspended the pellets in 50 ug/ml Calcoflour (Sigma), incubated for 5 minutes at room temperature, and washed an additional two times in PBS. We imaged cells with an Axiovert 200 M microscope (see below) and 0.2 um z-stack slices were taken. A representative image of the maximum projection for each genotype is shown.

### Microscopy and image preparation

We performed microscopy with an Axiovert 200 M microscope, a DeltaVision Core microscope or a Spinning Disk Confocal system as indicated below. We took images in Figure 4A, S1A, S1B, S4B, S4C and S5A on an Axiovert 200 M microscope (Carl Zeiss MicroImaging, Inc.) fit with a 100×/1.45-numerical aperture oil immersion objective and Cascade II-512B camera (PhotoMetrics, Inc.). Images were acquired using Openlab software (v5.5.0; Improvision), and FIJI and Photoshop (Adobe) were used to make linear adjustments to brightness and contrast. We used the DeltaVision Core fit with a U PLAN S APO 100×, 1.4 NA objective (Olympus) and a CoolSnapHQ2 Camera (Photometrics) for images in Figure 5E. A z-series of 0.2 um step size was taken at the indicated time after release from mitotic arrest. Images were deconvolved using softWoRx’s (Applied Precision Ltd.) iterative, constrained three-dimensional deconvolution method. FIJI was used to make linear adjustments to brightness and contrast. A representative single slice image is shown at each time. We used the Spinning Disk Confocal system (Leica) fit with a CSU-X1 spinning disk head (Yokogawa Electric Corporation), a PLAN APO 100×, 1.44 NA objective (Leica), and an Evolve 512 Delta EMCCD camera (Photometrics) for the images in Figure 5A and 6E. A step-size of 0.2 um was used and images were acquired using Metamorph (Molecular Devices) and deconvolved using AutoQuant X3’s (Media Cybernetics) iterative, constrained three-dimensional deconvolution method. FIJI was used to make linear adjustments to brightness and contrast. A representative maximum projection image is shown.

## Acknowledgments

We thank members of the Weiss and L. Lackner labs at Northwestern University for critical review of the manuscript. We thank K. Labib, G. Pereria, F. Cross, E. Bi, R. Lamb, Y. Barral, and S. Strahl for strains, plasmids and antibodies. We thank C. Wilke (Northwestern University Biological Imaging Facility) for assistance with electron microscopy, and E. Anderson and S. Zdraljevic for assistance with R software (statistical analysis and graphing). We also thank staff and instrumentation support from the Biological Imaging Facility and High Throughput Analysis Laboratory at Northwestern.

## Author Contributions

Conceptualization, J.L.B and E.L.W; Methodology, J.L.B and E.L.W; Investigation, J.L.B, M.D. and E.L.W; Writing, J.L.B and E.L.W; Visualization, J.L.B and E.L.W; Funding Acquisition, E.L.W.

## List of Abbreviations

AMR: Actomyosin ring
PS: Primary septum
SS: Secondary septum
CDK: Cyclin dependent kinase
AID: Auxin inducible degron
IP: Immunoprecipitation
MEN: Mitotic Exit Network
RAM: Regulation of Ace2 and Morphogenesis network

